# Deciphering the Regulatory Landscape of *γδ* T Cell Development by Single-Cell RNA-Sequencing

**DOI:** 10.1101/478529

**Authors:** Sagar, Maria Pokrovskil, Josip S. Herman, Shruti Naik, Elisabeth Sock, Ute Lausch, Michael Wegner, Yakup Tanriver, Dan R. Littman, Dominic Grün

## Abstract

Recent studies have established *γδ* T cells as critical players in a broad range of infections, antitumor surveillance, autoimmune diseases and tissue homeostasis. However, differentiation of *γδ* T cells in the adult thymus remains poorly understood, due to the rare frequency of this lineage. Here, we infer high-resolution developmental trajectories of this rare population by single-cell RNA-sequencing. We reveal previously unknown subtypes and identify the transcription factor c-MAF as a novel key regulator of IL-17-producing *γδ* T cell (*γδ*T17) differentiation. c-MAF knockout mice exhibit a complete block in *γδ*T17 differentiation, absence of these cells from peripheral organs, and protection from an autoimmune phenotype in a psoriasis model. Single-cell RNA-sequencing of *Sox13* and *Rorc* knockout mice pinpoints c-MAF as an essential missing link between these lineage-specifying factors. These findings significantly enhance our understanding of *γδ* T cell ontogeny. Our experimental strategy provides a blueprint for deciphering differentiation of rare cell types.

## INTRODUCTION

Traditionally, the central concepts pertaining to the immune system have strongly demarcated innate and adaptive branches of immunity until the discovery of relatively new cell types which seem to blur this binary classification (Lanier, 2013). *γδ* T cells represent one such lineage: Although displaying somatically rearranged antigen receptors on their cell surface, akin to adaptive αβ T cells and B cells, several fundamental differences in their development and functions make them traverse the strict conventional definitions of innate and adaptive immunity (Vantourout and Hayday, 2013).

*γδ* T cells are the earliest T-cells to develop in all vertebrate species examined to date, and mainly populate barrier epithelial tissues such as skin, lung, intestine and reproductive tract. Moreover, their differentiation is developmentally pre-programmed: Rearrangement of defined T cell receptor (TCR) *γ*-chains occurs at discrete time points and is followed by selective migration of the cells to individual epithelial tissues during fetal development. *γδ* T cell development continues after birth albeit utilizing different TCR *γ*-chains (Carding and Egan, 2002). Functionally, *γδ* T cells exert innate-like rapid immune responses by recognizing a broad spectrum of molecules including non-peptide antigens through TCR-dependent and -independent mechanisms (Hayday, 2009). Their effector functions are pre-programmed in the thymus prior to egress. Therefore, when activated, *γδ* T cells can rapidly produce various cytotoxic molecules, such as perforin and granzymes as well as effector cytokines, such as IL-17, IFN-γ and IL-4, and regulate an early innate response by recruiting neutrophils during infection, hypersensitivity and autoimmunity. Moreover, they have been shown to play immunosuppressive roles during injury resolution and to release growth factors leading to epithelial tissue repair and wound healing (Bonneville et al., 2010).

Although the role of *γδ* T cells in various aspects of infection, tumor and autoimmune diseases is widely recognized, their intrathymic differentiation is not well understood, in contrast to *αβ* T cell differentiation (Munoz-Ruiz et al., 2017). The reason for this lack of knowledge is partly due to the fact that, in the adult thymus, *γδ* T cell subsets are rare in comparison to conventional *αβ* T cells. Furthermore, the transcriptional identity of different subtypes developing in the thymus is not characterized at a high-resolution. Current studies favor a TCR-dependent two-step model of *γδ* T cell commitment and subsequent effector differentiation. TCR signal strength has been shown to play an essential role in *γδ* T cell commitment where weak signals promote *αβ* commitment and strong signals promote the *γδ* T cell fate. Furthermore, the subsequent effector differentiation into IL-17, IFN-*γ* and IL-4 producing *γδ* lineages also require varying levels of TCR signals. Weak signals promote IL-17 producing *γδ* T cell (*γδ*T17) lineage whereas progressively stronger signals promote IFN-*γ* and IL-4 producing *γδ* lineages, respectively (Fahl et al., 2014; Zarin et al., 2015). The underlying molecular mechanisms governing this process still remain enigmatic despite years of intense research. Moreover, the regulatory hierarchy controlling the emergence of distinct sub-types downstream of TCR signaling strength-dependent *γδ*-lineage commitment is almost entirely uncharacterized. Especially, how a high TCR signal-experiencing *γδ*-committed progenitor downregulates these signals to differentiate into *γδ*T17 lineage is an intriguing open question.

In order to gain insights into these processes and decipher the underlying regulatory mechanisms, we utilized multi-color flow cytometry to enrich for rare cell populations encompassing all differentiation stages of the *γδ* T cell lineage in the adult murine thymus and profiled them using single-cell RNA-sequencing (scRNA-seq). From these data, we predict continuous differentiation trajectories by pseudo-time analysis and infer gene regulatory network modules governing each stage of differentiation. We further identified a number of novel *γδ* sub-types or -states. This analysis reveals c-MAF as a previously unknown critical regulator of *γδ*T17 development. We investigated functional consequences of the loss of c-MAF in the *γδ* T cell lineage and observed a lack of ROR*γ*t^+^ *γδ* T cells in peripheral tissues as well as resistance to imiquimod-induced psoriasis-like skin inflammation in *Maf*-deficient mice. To decipher the regulatory hierarchy of c-MAF, encoded by *Maf*, and the two *γδ*T17 lineage-determining factors *Sox13* and *Rorc*, we performed a comparative scRNA-seq analysis of thymocytes from knockout mice of these three factors. Our data not only provide a high-resolution transcriptome-wide map of differentiation of this rare T cell lineage but also functionally validate the de novo predicted regulatory cascade of *γδ*T17 differentiation including *Sox13*, *Maf* and *Rorc*. Collectively, our study demonstrates the power of combining enrichment of rare populations by flow cytometry with deep single-cell RNA-seq to comprehensively sample differentiation trajectories of rare cell types. We showcase that computational regulatory network inference can be extended and validated by a merged analysis of wild-type and knockout animals for central predicted regulators, in order to take scRNA-seq experiments beyond the mere descriptive level.

## RESULTS

### scRNA-seq of rare cell types isolated by flow cytometry recovers all stages of T cell commitment and *γδ* T cell differentiation

To investigate the transcriptional landscape of *γδ* T cell differentiation, we isolated thymocyte subsets from 6-weeks old female mice utilizing established cell surface markers in order to enrich for the rare thymic progenitors as well as cell types of the *γδ* T cell lineage (Figure S1A). These populations comprise rare bipotent *αβ*/*γδ* T cell precursors – double negative (DN) 1, DN2 and DN3 (Figure S1B), CD25^+^ earliest rare *γδ* T cell precursors, previously defined as pre-selected and post-selected *γδ* T cells (Prinz et al., 2006) (Figure S1C), pan *γδ* T cells (mainly containing CD24^+^ immature population) and CD24^-^ mature population (Figure S1D), as well as IFN-*γ* producing CD122^+^ *γδ* T cells (Narayan et al., 2012; Shibata et al., 2008) (Figure S1E). Using this strategy, we expected to sample the entire *γδ* T cell differentiation trajectory. In total, cells were isolated from 11 female mice with each population purified from at least two animals. Single cells were sorted in 384-well plates by flow cytometry to perform scRNA-seq according to our published mCEL-Seq2 protocol (Hashimshony et al., 2016; Herman et al., 2018) and downstream data analyses were conducted using the RaceID3 algorithm (Figure 1A). After removing low-quality cells based on low total transcript numbers, 4146 cells were included in the final analysis. We observed no batch-associated variability in the dataset and recovered all cell types from different replicates. RaceID3 identified 31 clusters comprising >5 cells demonstrating substantial heterogeneity within conventionally defined populations of the T cell progenitors and the *γδ* T cell lineage (Figures 1B and 1C). First, we investigated the expression of known regulators of T-cell commitment (e.g., *Gata3*, *Runx1*, *Tcf7* and *Bcl11b*) as well as transcription factors involved in *γδ* T cell differentiation such as *Sox13*, *Rorc* (encoding ROR*γ*t), *Tbx21* (encoding T-BET) and *Eomes*. We were able to detect the expression of these genes in our dataset and the expression pattern confirmed the robustness of our cell clustering approach, using RaceID3 (Figure S1F). The level of heterogeneity observed in each of the conventionally defined progenitor populations suggests the existence of previously unknown sub-types or - states. In summary, combining multi-color fluorescence-activated cell sorting (FACS) to enrich for rare cell types with scRNA-seq is a powerful strategy to study differentiation of rare cell types. Using this strategy, we recovered all cell types during T cell commitment and *γδ* T cell differentiation.

**Figure 1.**
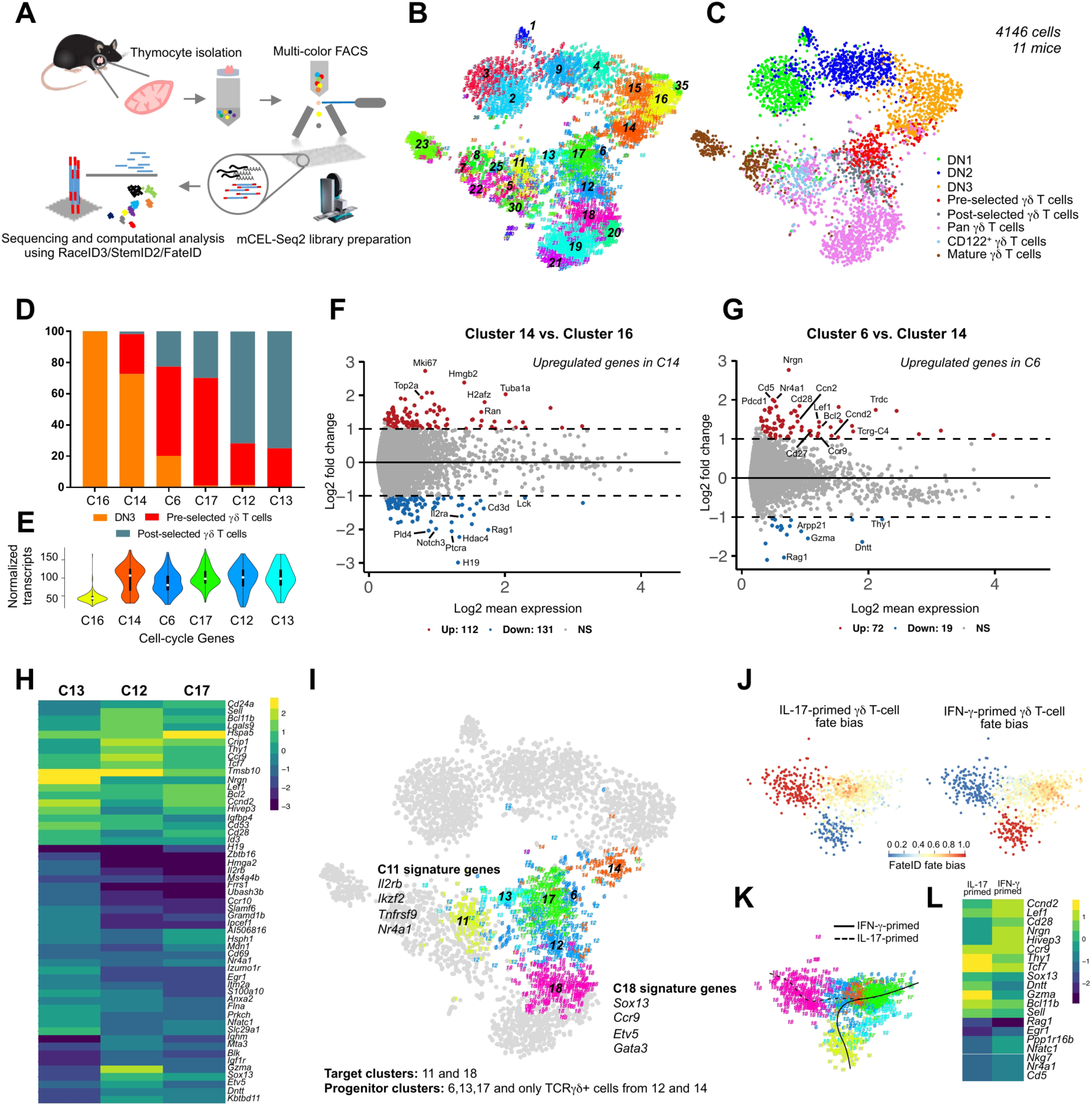
Single-cell RNA-sequencing (scRNA-seq) of cells encompassing all stages of *γδ* T cell differentiation. **(A)**Schematic representation of the workflow used for single cell sorting, library preparation and data analysis. **(B)** t-SNE representation based on transcriptome similarities showing the clusters identified by RaceID3 algorithm. 31 clusters comprising >5 cells are depicted in black color. **(C)**t-SNE representation showing the sorted cell types. Colors represent different cell types sorted using fluorescence-activated cell sorting (FACS) (n=11, 6-week old female mice).**(D)**Proportion of DN3, pre-selected and post-selected *γδ* T cells in clusters 16, 14, 6, 17, 12 and 13. Note the gradual increase in *γδ* T cell progenitors. **(E)**Violin plot showing the aggregated normalized transcript counts of cell cycle-related genes in these clusters. **(F)**MA plot showing differentially expressed genes between clusters 14 and 16. Cell cycle-related genes were upregulated in cluster 14. **(G)**MA plot showing differentially expressed genes between clusters 6 and 14. Cluster 6 exhibited the upregulation of TCR signal strength-related genes such as *Nr4a1* and *Cd5*. **(H)**Heatmap showing the differentially expressed genes in clusters 13, 12 and 17. Shortlisted genes showed a minimum of 2-fold change in at least one of the pairwise cluster comparisons (adjusted *P* < 0.05). **(I)**t-SNE representation showing the clusters included in the fate bias analysis using the FateID algorithm. Clusters 11 and 18 were selected as target clusters and the fate bias was calculated in clusters 6, 13, 17 as well as *γδ* progenitors in clusters 12 and 14. **(J)** Classical multidimensional scaling (CMD) dimensional reduction representation of all the clusters included in the fate bias analysis. Color scale represents fate bias probabilities on the scale of 0 to 1. **(K)** CMD representation showing the principal curves fitted to cells for each of the two lineages. **(L)** Heatmap showing the few shortlisted differentially expressed genes between progenitors with a fate bias >0.5 towards either of the two lineages (adjusted *P* < 0.05).

### Characterizing transcriptional and cellular heterogeneity in the early double negative T cell progenitors

We first characterized the heterogeneity in the DN1-DN3 progenitors capable of giving rise to both *αβ* and *γδ* T cell lineages. DN1 cells, also known as early thymic progenitors (ETPs), possess the potential to give rise to alternate hematopoietic lineages. ETPs are known to express genes associated with proliferation and leukemia (Yui and Rothenberg, 2014). Accordingly, we detected the expression of genes such as *Bcl11a*, *Lmo2*, *Hoxa9*, *Lyl1*, *Hhex* and *Bmyc* in ETPs (Figures S2A and S2D). The expression of *Flt3*, which has been associated with the most primitive ETPs entering the thymus, was restricted to a subset of ETPs (Figure S2D). RaceID3 classified ETPs into two distinct clusters – clusters 2 and 3 (Figures 1B and 1C). A closer examination of these clusters revealed that cluster 3 exhibits higher expression of receptor tyrosine kinase *Kit* and the mast cell associated gene *Cpa3*, while cluster 2 expresses higher levels of genes associated with DNA replication and cell cycle progression such as minichromosome maintenance genes (e.g., *Mcm2*, *Mcm5* and *Mcm6*) as well as *Pcna*, *Ran* and *Ccnd2* (Figure S2B). In conclusion, based on single cell transcriptome analysis, we identified two ETP subsets – one subset expressing multipotency markers and another undergoing cell cycle progression.

T cell commitment happens at the DN2 stage. DN2 cells are further subdivided into the uncommitted DN2a and committed DN2b subtypes, a transition marked by the downregulation of *Kit* and upregulation of T cell commitment factor, *Bcl11b* (Kueh et al., 2016; Yui et al., 2010). Noticeably, although we sorted DN2 cells in an unbiased fashion, our clustering approach identified two major subsets of DN2 progenitors – clusters 4 and 9 (Figures 1B and 1C). Cluster 9 resembled the DN2a subset as it exhibited an ETP-like gene expression signature (e.g., *Lyl1*, *Spi1*, *Adgrg3*). Cluster 4 showed an upregulation of *Bcl11b* as well as T cell specific genes such as *Cd3g*, *Cd3e*, *Cd3d*, *Lck*, *Lat* and *Notch1* and thus corresponds to a DN2b-like committed T cell progenitor cluster (Figure S2B).

In the T cell committed DN3 compartment, we identified four different clusters – clusters 14-16 and 35 (Figures 1B and 1C). Clusters 15, 16 and 35 represent cells undergoing recombination and hence express *Rag1* and *Rag2* (Figure S1D) and pre-T cell antigen receptor alpha, *Ptcra* but have minimal levels of cell cycle-related genes (Figures S2C and S2D). Cluster 14 comprises proliferating cells expressing *Mki67* and *Pcna* and may represent beta-selected DN3 cells (Figures S2B and S2C). Overall, the DN3 compartment exhibits substantial heterogeneity, consisting of cells underdoing recombination, selection and subsequent proliferation. Collectively, our analysis revealed a continuous transcriptional shift in early T cell progenitors undergoing commitment, recombination and selection tightly coupled with proliferation.

### TCR signal strength-related differentiation propensity among CD25^+^ *γδ* T cell progenitors

Next, we sought to characterize the transcriptional signature of the early *γδ* T cell progenitors. Since it has been shown that during the DN2-DN3 transition, bipotent *αβ*/*γδ* T cell precursors become lineage restricted, we sequenced rare pre-selected and post-selected *γδ* T cell progenitors (~0.005% of total thymocytes) from the DN3 compartment (Ciofani et al., 2006; Prinz et al., 2006). These cells were grouped into five different clusters, i.e. clusters 6, 12, 13, 14 and 17 (Figures 1B and 1C). Few of the pre-selected *γδ* T cell progenitors co-clustered with DN3 cells (Cluster 14) and express the proliferation marker *Mki67*, but no *γδ*-specific signature genes (Figures 1D-1F). *γδ* T cell progenitors in cluster 6 express TCR signaling strength-related genes such as *Nr4a1* (encoding NUR77) and *Cd5* (Figure 1G). Furthermore, they express *Cd28*, *Pdcd1* and *Lef1* as well as the *γδ*-specific transcription factor, *Sox13* (Figures 1G and 2C). Interestingly, cells within cluster 12 exhibit increased expression of *Sox13* and *Blk* (Figures 1H and 2C). *Blk* knockout mice show impaired *γδ*T17 cell differentiation and augmented TCR signaling (Laird et al., 2010). Therefore, this indicates that cells comprising cluster 12 are already primed towards the *γδ*T17 differentiation pathway whereas cells in cluster 13 downregulate *Sox13* and express TCR signal strength-related genes, such as *Nr4a1* and *Egr1*, at higher levels (Figure 1H). Cluster 13 cells also express *Nfatc1* and protein kinase C family member, *Prkch* indicating that these cells may experience higher TCR signals (Figure 1H). Altogether, we identified CD25^+^ *γδ* T cell subsets having distinct gene signatures in terms of proliferation, TCR signals and *γδ* T cell commitment.

**Figure 2.**
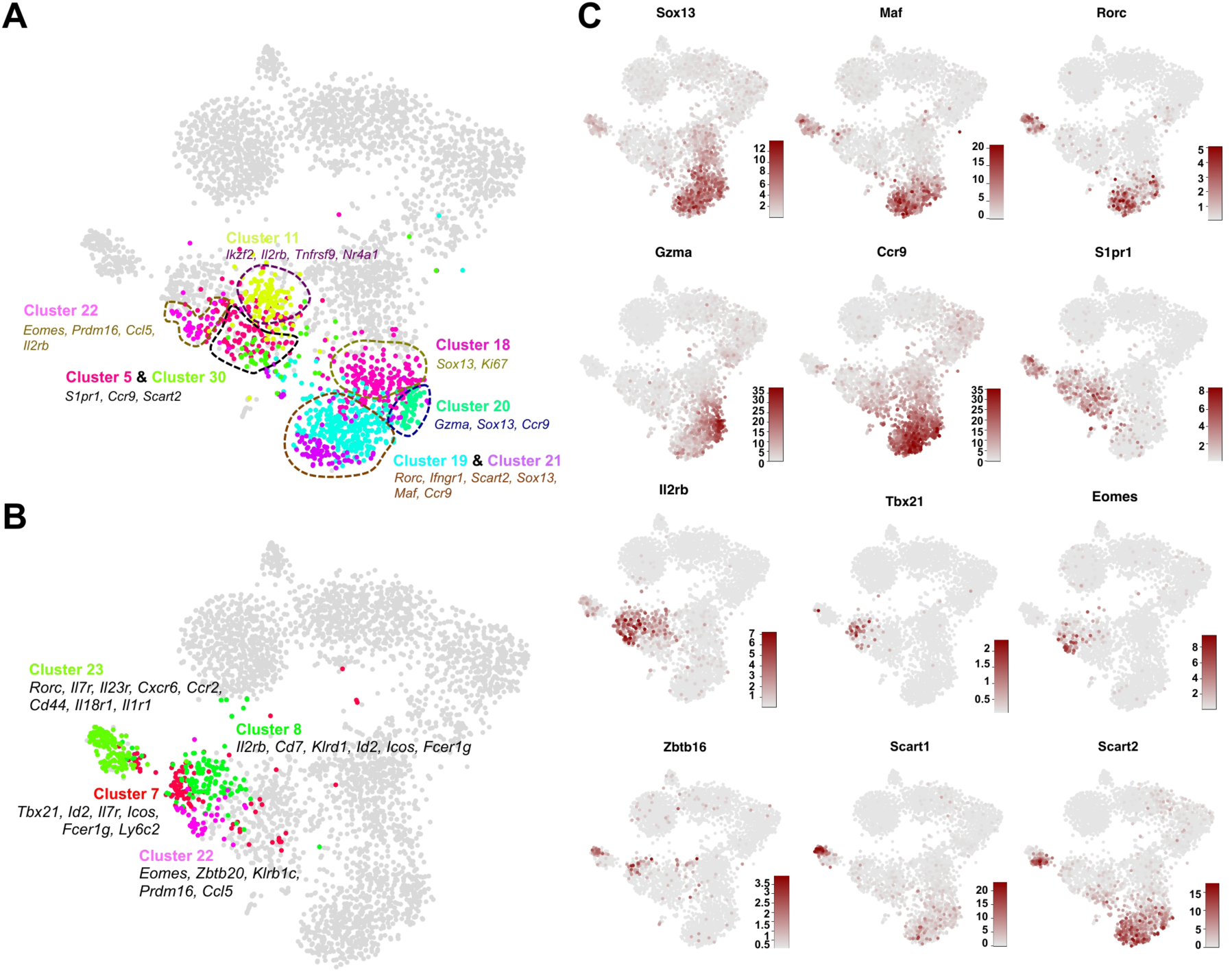
Immature and mature *γδ* T cells consist of various cell subtypes and exhibit substantial transcriptional heterogeneity. **(A)** t-SNE representation highlighting the clusters enriched in immature *γδ* T cells. Other cells are shown in grey. We identified six different subtypes. Clusters are circled and labeled. Few marker genes characterizing these clusters are also depicted. **(B)** t-SNE representation showing the clusters enriched in mature *γδ* T cells. Clusters are labeled and few marker genes characterizing these clusters are also highlighted. **(C)** t-SNE representation showing the expression of key marker genes differentially expressed among various *γδ* cell subtypes. Color scale represents the normalized transcript counts.

To further validate the existence of cell types experiencing different levels of TCR signaling in the pre-selected and post-selected *γδ* T cell compartment, we utilized FateID, our recently developed algorithm for the quantification of fate bias in the progenitor compartment towards the distinct mature lineages. FateID utilizes the transcriptional signature of mature lineages, termed target clusters, on the basis of which the fate probabilities of each progenitor cell are inferred. Since cluster 11 and cluster 18 represent the earliest *γδ*T17-primed and IFN-*γ* primed immature *γδ* T cell types, respectively, we quantified the fate bias among the CD25^+^*γδ* T cell progenitors using clusters 11 and 18 as target clusters (Figure 1I). The FateID-derived lineage probabilities clearly discriminated cells with a preferential bias towards the *γδ*T17 or IFN-*γ* producing lineage, respectively (Figures 1J and 1K). Differential gene expression analysis between cells biased towards the distinct lineages (fate bias probability >0.5) revealed that *γδ*T17 lineage-primed cells expressed higher levels of genes such as *Bcl11b*, *Thy1*, *Ccr9* and *Sox13* whereas IFN-*γ* lineage-primed cells expressed high TCR signal strength-related genes – *Cd28*, *Nfatc1*, *Nr4a1* and *Egr1* (Figure 1L). Taken together, our data identified TCR signal strength-related transcriptional priming in the early CD25^+^ *γδ* T cell progenitors and support the TCR signal strength-based effector *γδ* T cell differentiation model (Zarin et al., 2015).

### Identifying transcriptionally heterogeneous immature and mature *γδ* T cell subsets

Thirty years after the discovery of *γδ* T cells, the full extent of the diversity of *γδ* T cell subsets remains elusive. Since they rearrange different antigen receptors, reside in different tissues and perform different biological functions, a substantial heterogeneity of sub-types already during intrathymic development is expected. As scRNA-seq is the method of choice to reveal transcriptionally different *γδ* T cell subsets, we analyzed immature and mature *γδ* T cells in our dataset.

We found six different subsets in the CD24^+^ immature *γδ* T cell compartment (Figure 2A). As suggested by the gene expression signature, cluster 18 (expressing *Sox13* and *Mki67*) represents a progenitor cluster, giving rise to clusters 19 and 20 (Figures 2A and 2C). Cluster 20 expresses very high levels of granzyme A (*Gzma*) representing a newly identified *γδ* T cell subtype (Figure 2C). This cluster may portray a novel cytotoxic subpopulation of *γδ* T cells. Noticeably, some of *Gzma*^hi^ cells expressed *Rorc*. Cluster 19 potentially represents an intermediate cell type, giving rise to the *Rorc*^+^ cluster 21 (Figure 2A), since expression of *Rorc* begins in cluster 19 and peaks in cluster 21 (Figure 2C). Based on the expression of *Sox13*, *Rorc* and *Ccr9*, cluster 21 can be considered the classical *γδ*T17 subpopulation (Figure 2C). We did not detect the expression of *Il17a* and *Il17f* in *Rorc*^+^ cells in cluster 21. Clusters 11 and 22 express *Il2rb* (encoding CD122) and comprise sorted CD122^+^ immature *γδ* T cells thereby representing the IFN-*γ* producing *γδ* T cells (Figure 2C). Cluster 22 further expresses *Eomes*, *Prdm16*, and *Ccl5* (Figure 2C). Few cells in this cluster also express *Klrb1c* (encoding NK1.1). Moreover, the *Eomes*^+^ cluster 22 contains cells from the immature as well as the mature *γδ* T cell compartment (Figures 1C and 2A). *Eomes* expression on *γδ* T cells is associated with the IFN-*γ* producing Th1-like subset, although it has been shown to be dispensable for IFN-*γ* production (Barros-Martins et al., 2016; Lino et al., 2017). Since *Eomes*^+^ cells represent a rare *γδ* T cell subset having a distinct transcriptional signature, it is likely that they serve unique characteristic functions during an immune response. Interestingly, we identified another uncharacterized subtype, represented by clusters 5 and 30, with no clear transcriptional signature (Figure 2A). These clusters are best discriminated by the expression of the thymocyte developmental genes *Ccr9* and *S1pr1,* which are crucial for egress from the thymus and migration to peripheral sites (Figure 2C). Except for these two clusters, the expression domains of *Ccr9* and *S1pr1* were mutually exclusive and found to be more restricted to *Sox13*^+^ and *Il2rb*^+^ *γδ* T cells, respectively. Although in the t-SNE (t-distributed stochastic neighbor embedding) representation, clusters 5 and 30 seem to be transcriptionally similar to *Il2rb*^+^ *γδ* T cells, they also expressed *Sox13* and *Scart2*, genes associated with *γδ*T17 differentiation, albeit at lower levels. It is thus likely that these cells ether exit the thymus in a potentially naïve state and are primed to their effector fates in the periphery or may represent the IL-17^+^ IFN-*γ*^+^ double producers’ progenitors (Serre and Silva Santos, 2013). Overall, we identified six transcriptionally unique immature *γδ* subsets of which two were previously uncharacterized.

We then focused on characterizing the transcriptional heterogeneity in the mature CD24^-^ *γδ* T cells. Broadly, mature *γδ* T cells were found to express two different transcription factors – *Tbx21* (clusters 7 and 8) and *Rorc* (cluster 23) while few cells also express *Eomes* (cluster 22) (Figures 2B and 2C). *Tbx21* is necessary for the differentiation of IFN-*γ* producing *γδ* T cells (Barros-Martins et al., 2016). We observed substantial heterogeneity in *Tbx21*^+^ and *Eomes*^+^ compartments (not resolved by clustering), based on the expression of various genes such as – *Icos*, *Ly6c2*, Ccl5, *Klrb1c*, *Klrd1* and *Zbtb16* (encoding PLZF) (Figure S3). Cluster 23 comprised *Rorc*^+^ cells and consisted of two major subpopulations (not resolved by clustering) demarcated by the expression of two scavenger receptors – *Scart1* and *Scart2* (Figures 2B and 2C). *Scart1*^+^ cells expressed *Zbtb16*, required for the differentiation of V*γ*6^+^*γδ*T17 cells developed in the fetus (Lu et al., 2015) (Figures 2C), suggesting that these cells are either recirculating V*γ*6^+^ *γδ*T17 cells or represent a V*γ*6^+^ thymic potential progenitor pool set aside during embryonic development. Of note, the expression of *Id2* and *Il7r* was upregulated in all the *γδ* subtypes transitioning from the immature to the mature state (Figure S3). Taken together, scRNA-seq revealed substantial heterogeneity of mature *γδ* T cells in the adult thymus.

In order to understand the relationship between the specific variable chains expressed by *γδ* T cells and their transcriptome, we profiled *γδ* T cells expressing different variable *γ* and δ chains on their cell surface (V*γ*1, V*γ*4 and V*δ*6.3) (Heilig and Tonegawa, 1986) (Figures S4A and S4B). In the immature *γδ* T cell compartment, as reported previously (Narayan et al., 2012), V*γ*4^+^ cells mainly co-clustered with the *γδ*T17 lineage while V*γ*1^+^ cells co-clustered with IFN-*γ* producing lineage (Figure S4C). Interestingly, both of the V*γ*1^+^ and V*γ*4^+^ *γδ* T cells co-clustered with *Gzma*^hi^ cells, suggesting that these cells can rearrange either of these TCR*γ* chains (Figure S4C). *Scart1*^+^ *Zbtb16*^+^ mature *γδ* T cells did not co-cluster with mature V*γ*4^+^ *γδ* T cells strengthening our assumption that these cells are fetal-derived V*γ*6^+^ *γδ* T cells (Figure S4C).

Intrathymic *γδ* T cell development starts during fetal development. These fetal-derived *γδ* T cells are the first T cells to be generated and colonize peripheral mucosal tissues (Carding and Egan, 2002). In order to compare *γδ* T cell differentiation in the embryo and adult, we further performed scRNA-seq of CD24^+^ and CD24^-^ *γδ* T cells from embryonic day 17.5 (E17.5) thymi and found that the majority of CD24^+^ *γδ* T cells were *Sox13*^+^ (Figures S5A-S5C). We were able to identify transcriptionally similar counterparts to specific week 6 subpopulations, i.e. *Rorc*^+^ *γδ*T17 cells (clusters 3 and 11), *Gzma*^hi^ subpopulation (cluster 8) as well as *Ccr9* and *S1pr1* double-positive cells (no specific cluster) (Figures S5A-S5C). Unlike their adult counterparts, embryonic *Rorc*^+^ cells present in clusters 3 and 11 expressed *Il17a* and *Il17f* (Figure S5D). Overall, as expected, E17.5 *γδ* T cells were more proliferative than week 6 cells expressing *Mki67* (Figure S5C). Strikingly, unlike week 6 *γδ* T cells, *Sox13*^+^ E17.5 *γδ* T cells expressed substantially higher levels of *Nr4a1* (Figure S5C), hinting towards a different requirement of TCR signals during embryonic *γδ* T cell differentiation. In the E17.5 thymi, mature CD24^-^ *γδ* T cells mainly expressed *Il2rb* but rarely *Tbx21* and *Eomes* (data not shown). The majority of these cells expressed *Klrd1* (encoding CD94) and *Nt5e* (encoding CD73) while few cells also expressed *Gzmb* and *Klrb1c* (Figures S5B and S5C). In conclusion, embryonic *γδ* T cells consist of several subsets similar to their adult counterparts but also exhibit particular transcriptional differences in terms of differentiation, proliferation and TCR signal related genes.

### Inferring differentiation trajectories and gene regulatory networks regulating *γδ* T cell development

In order to understand the regulatory control of *γδ* T cell differentiation in the thymus, we sought to infer the differentiation trajectories of this process using our scRNA-seq data. We applied the StemID2 algorithm to infer a lineage tree de novo (Grun et al., 2016; Herman et al., 2018) (Figure 3A) and derived the pseudo-temporal gene expression changes during

**Figure 3.**
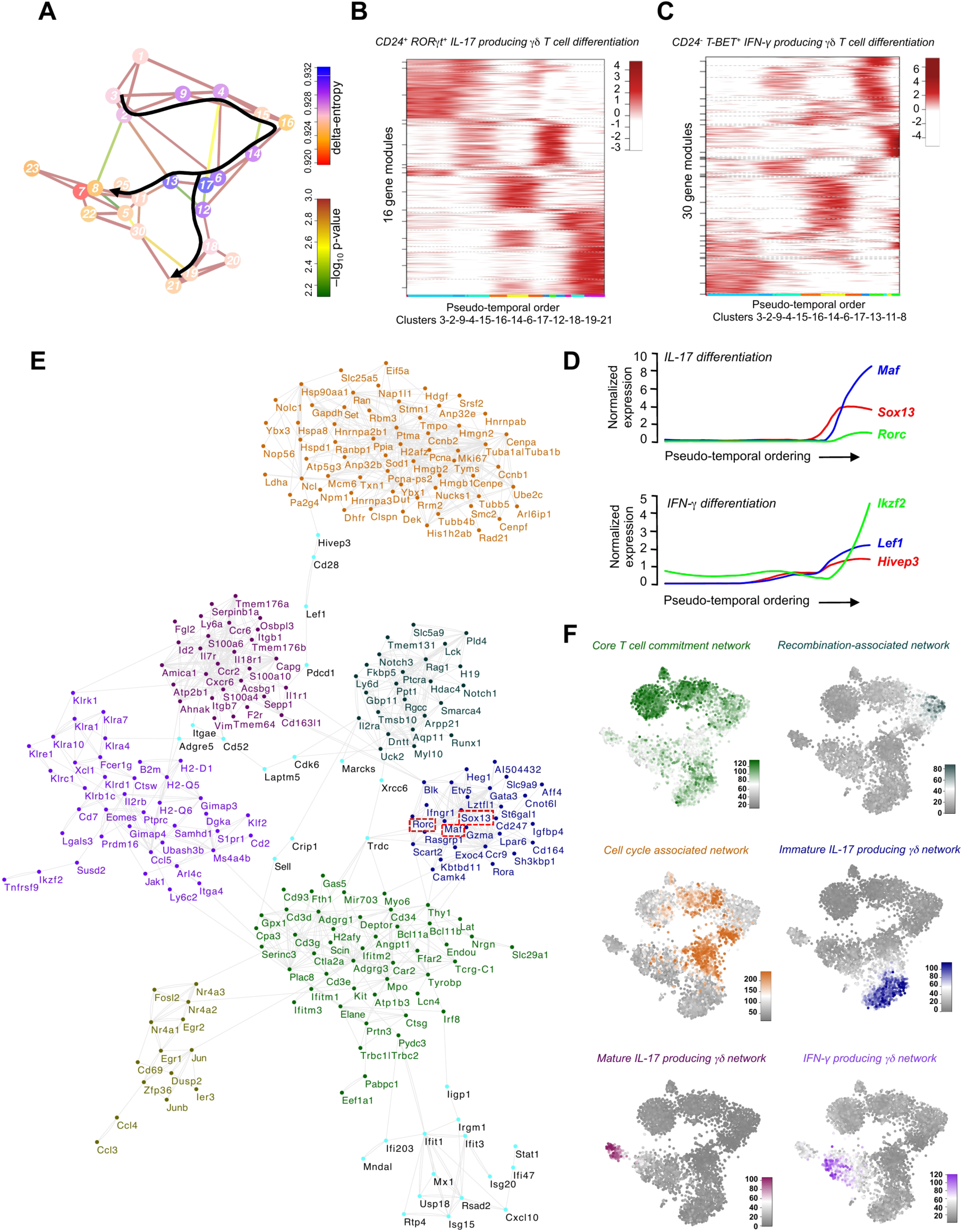
*γδ* differentiation trajectory analysis and gene regulatory network inference using single-cell RNA-sequencing data. **(A)** Inferred lineage tree of *γδ* T cell differentiation using StemID2 algorithm. Only significant links are shown (*P* < 0.01). The color of the link indicates the −log_10_*P*. The color of the vertices indicates the entropy. The thickness indicates the link score, reflecting how densely a link is covered with cells. The superimposed black arrows indicate the clusters used to generate pseudo-temporal profiles using self-organizing maps (SOMs). **(B)** SOM of z-score-transformed, pseudo-temporal expression profiles along the IL-17 producing *γδ* T cell (*γδ*T17) differentiation trajectory (clusters 3, 2, 9, 4, 15, 16, 14, 6, 17, 12, 18, 19 and 21). The color-coding at the bottom indicates the cluster of origin. The SOM identified 16 different modules of co-regulated genes. **(C)** SOM of z-score-transformed, pseudo-temporal expression profiles along the IFN-*γ* producing *γδ* T cell differentiation trajectory (clusters 3, 2, 9, 4, 15, 16, 14, 6, 17, 13, 11 and 8). The color-coding at the bottom indicates the cluster of origin. The SOM identified 30 different modules of co-regulated genes. **(D)** Plots showing the expression of *Sox13*, *Maf* and *Rorc* (top) as well as *Hivep3*, *Lef1* and *Ikzf2* (bottom) from the pseudo-temporal expression profiles along the IL-17 and IFN-*γ* producing *γδ* T cell differentiation trajectory, respectively. The lines indicate the pseudo-temporal expression values derived by a local regression of expression values across the ordered cells. **(E)** Gene regulatory network, as inferred from the single cell data of 6-week old thymi (**Figure 1**) using GENIE3 algorithm. The data of the top 1500 interactions were used to construct the gene regulatory network. Recovered network modules are highlighted in different colors. Note the presence of *Sox13*, *Maf* and *Rorc* (highlighted using dashed red rectangles) in one on the modules (blue). **(F)** t-SNE representation showing the aggregated expression of genes present in the different modules.

*γδ*T17 and IFN-*γ* producing *γδ* T cell differentiation using self-organizing maps (SOMs). SOMs of pseudo-temporal gene expression profiles along the predicted *γδ*T17 and IFN-*γ* producing differentiation trajectories grouped the expression profiles into 16 and 30 gene modules, respectively (Figures 3B and C). These different gene modules are activated at different time points during *γδ* T cell development. We then focused on the late modules activated during the effector differentiation of *γδ* T cells into *γδ*T17 and IFN-*γ* producing lineages. These late modules predicted the role of various genes including transcription factors with known and unknown functions during the differentiation of *γδ* lineages (Figures 3D and S6). Interestingly, histone-modifying factors were specifically upregulated during *γδ*T17 differentiation (Figure S6A). Furthermore, we inferred a gene regulatory network (GRN) from scRNA-seq data using the random forests-based GENIE3 algorithm (Huynh-Thu et al., 2010). We successfully recovered a GRN consisting of various gene modules (Figure 3E). A closer examination of these modules revealed that they characterize different stages of T cell commitment, TCR rearrangements, and *γδ* T cell development. We further plotted the aggregated expression of all genes comprising each gene module on the t-SNE map and found that different clusters, reflecting a distinct differentiation stage showed upregulated expression of specific gene modules indicating that GRN inference successfully recovered gene modules governing different stages of *γδ* T cell development (Figure 3F). In summary, we successfully mapped *γδ*T17 and IFN-*γ* producing differentiation trajectories using the scRNA-seq data and identified several regulatory factors involved in the process of *γδ* T cell development and effector differentiation.

### c-MAF is essential for *γδ*T17 differentiation in the embryonic and the adult thymus

Next, we were interested to understand the molecular mechanisms that lead to the suppression of TCR signaling and activation of the IL-17 program during *γδ*T17 differentiation. First, we analyzed the pseudo-temporal gene expression changes along the predicted *γδ*T17 differentiation trajectory. Interestingly, pseudo-temporal gene expression analsyis implicated an early role of *Sox13*, a subsequent role of *Maf* (encoding c-MAF) and a late role of *Rorc* in activating the *γδ*T17 effector program in *γδ* T cell progenitors (Figure 3D). Furthermore, gene regulatory network inference recovered an immature *γδ*T17-specific network module containing several genes including *Sox13*, *Maf* and *Rorc* (Figure 3E). Although the role of SOX13 and ROR*γ*t has been explored in *γδ*T17 differentiation (Barros-Martins et al., 2016; Gray et al., 2013; Ivanov et al., 2006; Malhotra et al., 2013; Melichar et al., 2007), the function of c-MAF in this differentiation process has not yet been investigated. c-MAF has been shown to regulate the differentiation of T helper 17, ROR*γ* t^+^ regulatory T cells, T follicular regulatory cells as well as IL-17 producing invariant natural killer T cells (Tanaka et al., 2014; Wheaton et al., 2017; Xu et al., 2009; Xu et al., 2018; Yu et al., 2017). In order to investigate the role of c-MAF in *γδ* T cell differentiation, we deleted *Maf* in lymphoid progenitors using *Il7ra^cre^*. *Maf* deletion did not affect intrathymic *αβ* T cell differentiation (Figures 6A and 6B). We then isolated and performed scRNA-seq of immature and mature *γδ* T cells from 6-week old *Maf^fl/fl^;Il7r^cre^* and *Maf^fl/fl^* thymi (Figures 4A and 4B). We found that immature *γδ* T cells from the knockout (KO) mice clustered separately from wild-type (WT) controls (Figure 4C). They failed to express *γδ*T17 signature genes such as *Rorc* and *Il17re* (Figure 4D). Differential gene expression analysis between the immature KO and WT *γδ* T cells revealed that *Maf*-deleted cells downregulated the expression of *Sox13*, *Sox4*, *Blk*, *Etv5*, *Bcl11b* and *Gata3* (Figure 4F). Importantly, *Maf*-deleted cells were found to upregulate the expression of TCR signal strength-related genes such as *Nr4a1* and *Cd69* (Figures 4E and 4F). Accordingly, gene set enrichment analysis between the immature KO and WT *γδ* T cells identified upregulation of AKT, MAP Kinase, Toll-like receptor (TLR) and receptor tyrosine kinase (RTK) signaling pathways in the *Maf*-deleted cells implicating an active role of c-MAF in downregulating TCR signaling components during *γδ*T17 differentiation (Figure 4G). Furthermore, *Maf*-deleted cells did not give rise to *Rorc*^+^ mature *γδ*T17 cells. All mature *Rorc*^+^ *γδ* T cell subtypes (*Zbtb16*^+^ and *Ztbtb16*^-^ populations) were missing in the KO thymi (Figure 4C). In conclusion, we identified c-MAF as a novel regulator required for the suppression of TCR signaling and the activation of the IL-17 program during *γδ*T17 differentiation.

**Figure 4.**
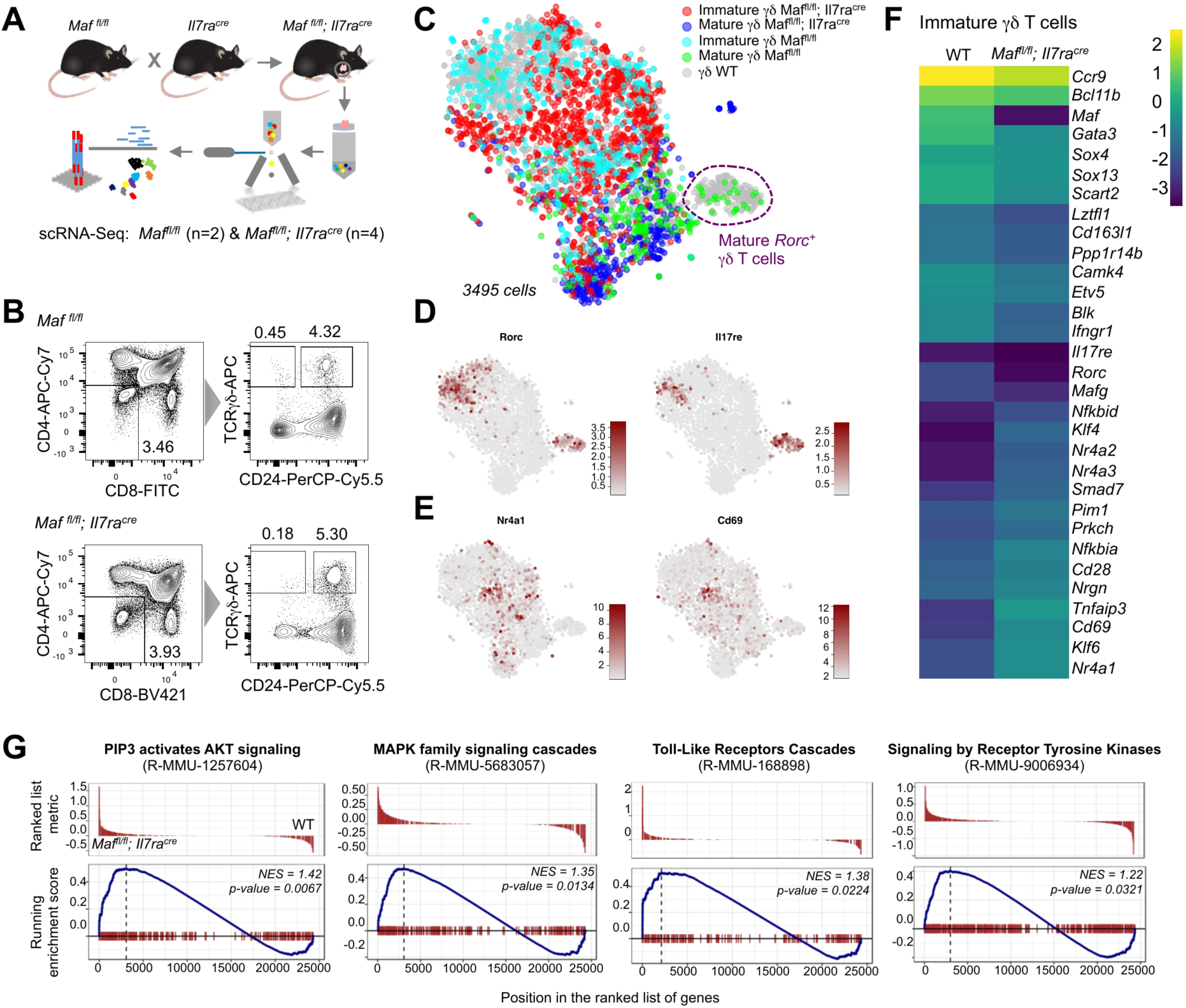
c-MAF is essential for the differentiation of *γδ*T17 lineage in the adult thymus. **(A)** Scheme showing the experimental design and scRNA-seq pipeline.**(B)**FACS plots depicting the sorting strategy for scRNA-seq experiments. Immature and mature *γδ* T cells were isolated from 6-week old *Maf^fl/fl^* and *Maf^fl/fl^;Il7r^cre^* female mice (n=2, wild-type (WT) and n=4, knockout (KO)). **(C)** t-SNE representation showing the sorted cell types. Note that the mature *Rorc*^+^ *γδ* T cell compartment lacks KO cells (outlined in purple). **(D)** t-SNE representation showing that the cells expressing *γδ*T17 signature genes *Rorc* and *Il17re* are present only in the WT mice. **(E)** t-SNE representation showing that few cells in the KO mice exhibited higher expression of TCR signal strength-related genes *Nr4a1* and *Cd69*. **(F)** Heatmap showing the differentially expressed genes between the CD24^+^ immature WT and KO *γδ* T cells. Shortlisted genes had adjusted *P* < 0.05. **(G)** Gene set enrichment analysis (GSEA) revealed that the AKT, MAP Kinase, Toll-like receptor and receptor tyrosine kinases pathways are enriched in *Maf*-deleted cells compared to the WT.

**Figure 5.**
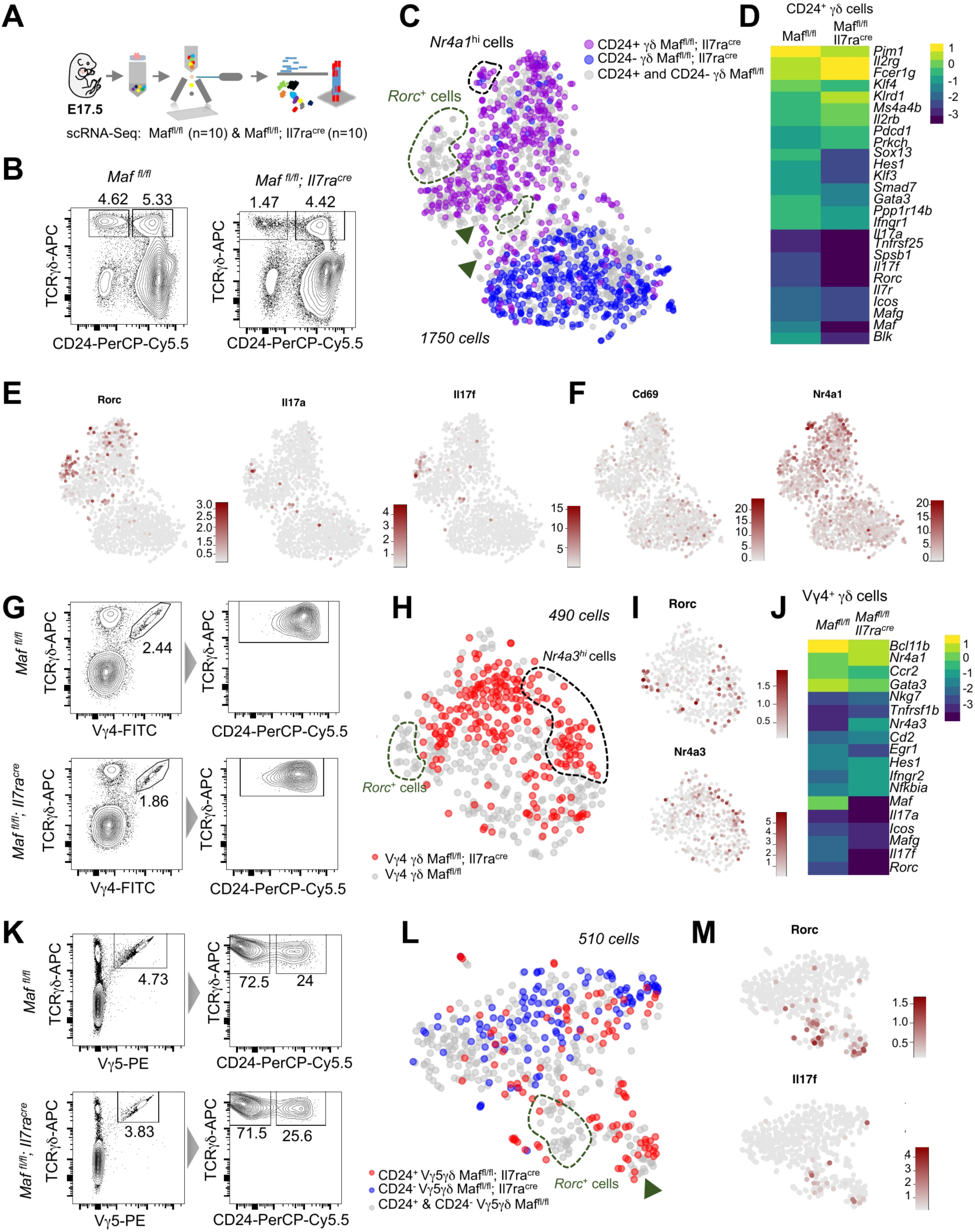
c-MAF is required for the development of *Rorc*^+^ *γδ* T cells in the embryonic thymus. **(A)**Scheme showing the experimental design and scRNA-seq pipeline.**(B)**FACS plots showing the sorted cell types. CD24^+^ and CD24^-^ *γδ* T cells were sorted from *Maf^fl/fl^* and *Maf^fl/fl^;Il7r^cre^* E17.5 thymi (n=10 embryos, each genotype).**(C)**t-SNE representation showing the sorted cell types from WT and KO E17.5 thymi. Note that *Rorc*^+^ cells are absent in the KO mice (green) and few cells exhibit higher *Nr4a1* and *Cd69* expression (outlined in black).**(D)**Heatmap showing the differentially expressed genes between CD24^+^ *γδ* T cells from the WT and KO mice (adjusted *P* < 0.05).**(E)**t-SNE expression maps showing that the cells expressing *Maf*, *Rorc* and *Il17f* are absent in the KO mice.**(F)**t-SNE representations showing the expression of *Nr4a1* and *Cd69*.**(G)**FACS strategy to sort V*γ*4^+^ cells from E17.5 thymi. Note that most of the V*γ*4^+^ cells are CD24^+^. Cells were sorted from the WT as well as KO E17.5 thymi.**(H)**t-SNE representation of cell types sorted as shown in (G). *Maf* deletion led to the absence of *Rorc*^+^**(I)**cells (outlined in green) and few of the KO cells expressed higher levels of *Nr4a3* (outlined in black).**(J)**t-SNE representation of the expression of *Rorc* and *Nr4a3* for cells sorted as shown in (G).**(K)**Heatmap showing the differentially expressed genes between V*γ*4^+^ *γδ* T cells from the WT and KO thymi (adjusted *P* < 0.05).**(L)**FACS strategy to sort V*γ*5^+^ cells from the WT and KO E17.5 thymi.**(M)**t-SNE representation of cell types sorted as shown in (K). *Maf* deletion led to the absence of *Rorc*^+^cells in V*γ*5^+^ *γδ* T cell compartment (green).**(N)**t-SNE map showing the expression of *Rorc* and *Il17f* of cell types sorted as shown in (K).

**Figure 6.**
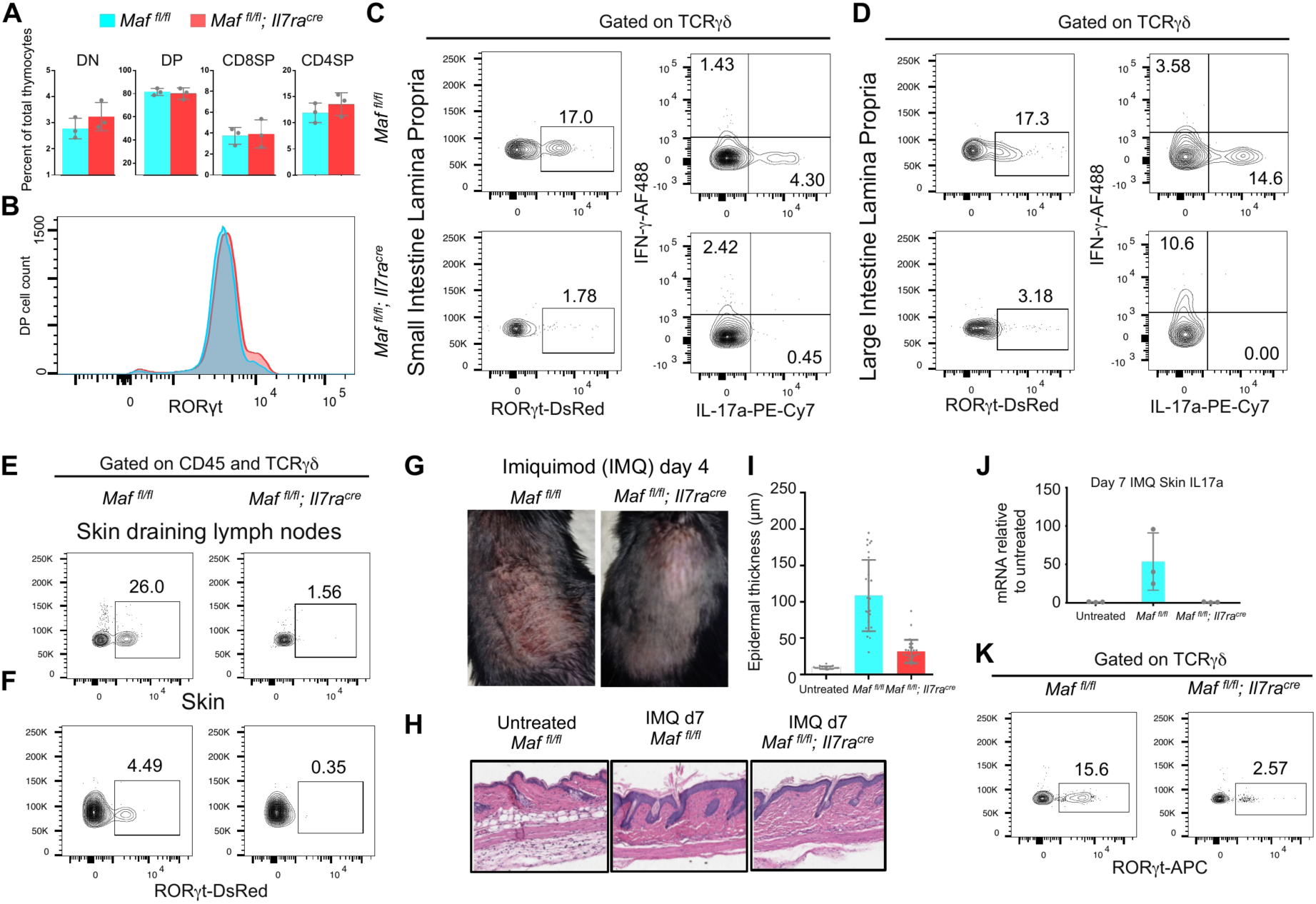
*Maf*-deficient mice lack peripheral ROR*γ*t^+^ *γδ*T17 cells and protected from psoriasis-like skin inflammation. **(A)** Percentage of double negative (DN), double positive (DP), CD8 and CD4 single positive (SP) cells in the 6-week old thymi from *Maf* KO and WT female mice (n=3, each genotype). **(B)**Intracellular staining showing the distribution of DP ROR*γ*t^+^ T cells in the 6-week old thymi from *Maf* KO and WT. **(C)**Left panel, Intracellular staining showing the quantification of ROR*γ*t^+^ *γδ* T cells from small intestine lamina propria (SILP) of 7-week old *Maf* KO and WT female mice. Right panel, Intracellular staining of IL-17a and IFN-*γ*after PMA and ionomycin stimulation, gated on TCR*γδ*^+^ SILP cells. **(D)**Left panel, Intracellular staining showing the quantification of ROR*γ* T^+^ *γδ* T cells from large intestine lamina propria (LILP) of 7-week old *Maf* KO and WT female mice. Right panel, Intracellular staining of IL-17a and IFN-*γ*after PMA and ionomycin stimulation, gated on TCR*γδ*^+^ LILP cells. **(E)**Fraction of ROR*γ* T^+^ *γδ* T cells in the skin-draining lymph nodes of 6-week old *Maf* KO and WT mice. **(F)**Fraction of ROR*γ* T^+^ *γδ* T cells in the skin of 6-week old *Maf* KO and WT mice. **(G)**Representative pictures of the dorsal skin of the 8-week old female *Maf* KO and WT mice after 4 days of consecutive IMQ application. **(H)**Hematoxylin and Eosin staining of the dorsal skin of untreated (WT) and treated (*Maf* KO and WT) mice. **(I)**Graph showing the quantification of epidermal thickness of untreated (WT) and treated (*Maf* KO and WT) mice. **(J)**RT-PCR showing the IL-17a mRNA levels in the skin of the *Maf* KO and WT mice after 6 consecutive days of IMQ treatment. **(K)**Intracellular staining showing the quantification of ROR*γ* T^+^ *γδ* T cells in the skin of the *Maf* KO and WT mice after 6 days of consecutive IMQ application.

Since *γδ*T17 cells colonizing the epithelial-rich tissues have been shown to be of embryonic origin (Haas et al., 2012), we further sequenced *γδ* T cells from KO and WT E17.5 thymi (Figures 5A and 5B). We found almost complete absence of *Rorc*^+^ *γδ* T cells in the E17.5 KO thymi (Figures 5C and 5E). Differential gene expression analysis between the KO and WT cells revealed that *Maf*-deleted cells did not express *Il17a* and *Il17f* (Figures 5D and 5E). As in the adult thymi, we also identified cell clusters expressing higher levels of *Nr4a1* and *Cd69* in the E17.5 KO *γδ* T cells (Figure 5F). Although unlike their adult counterparts, embryonic *Sox13*^+^ *γδ* T cells expressed *Nr4a1*, its expression was even higher in the embryonic KO *γδ* T cells compared to the WT. We also sequenced V*γ*4^+^ *γδ* T cells from the E17.5 KO and WT thymi (Figure 5G). *Rorc*^+^ cells were found to be missing in the *Maf*-deleted V*γ*4^+^ *γδ* T cell compartment (Figures 5H and 5I). Furthermore, these cells downregulated the expression of *Ccr2*, *Il17a*, *Il17f* and *Icos* while the expression of *Nr4a1*, *Nr4a3*, *Hes1*, *Cd2* and *Nfkbia* was higher in the KO cells (Figures 5I and 5J). In order to gain further insights into the role of c-MAF in the regulation of intrathymic differentiation of ROR*γ* T^+^ skin-resident *γδ* T cells, we specifically profiled V*γ*5^+^ *γδ* T cells from the E17.5 KO and WT thymi (Figure 5K). Indeed, scRNA-seq revealed that the V*γ*5^+^ *γδ* T cell compartment from KO thymi lacked *Rorc*^+^ *Il17f*^+^ *γδ* T cells (Figures 5L and 5M). Overall, our analysis clearly established that c-MAF is essential for the differentiation of all *Rorc*^+^ *γδ*T17 cell lineages in the embryonic as well as the adult thymi.

### Maf-deficient mice lack ROR*γ* T^+^ *γδ* T cells in peripheral tissues and are protected from psoriasis-like skin inflammation

To investigate the consequence of failed *γδ*T17 differentiation in the thymi of KO mice, we focused on characterizing the peripheral tissues in these animals. We observed a marked reduction in ROR*γ* T^+^ *γδ* T cells in the small and large intestinal lamina propria (Figures 6C and 6D). Consequently, the stimulation of *γδ* T cells with phorbol 12-myristate 13-acetate (PMA) and ionomycin isolated from these tissues revealed a drastic decrease in IL-17A-production (Figures 6C and 6D). We also confirmed the absence of ROR*γ* T^+^ *γδ* T cells in the skin draining lymph nodes (Figure 6E). The dermis of these mice also lacked ROR*γ* T^+^ *γδ* T cells (Figure 6F). Since *γδ*T17 cells have been shown to play an essential role in psoriasis-like skin inflammation (Cai et al., 2011), we investigated whether the absence of *γδ*T17 cells in *Maf* KO mice makes them resistant to psoriasis-inducing stimuli. Indeed, we found that the application of imiquimod (IMQ) on the dorsal skin, an established psoriasis model (Flutter and Nestle, 2013; van der Fits et al., 2009), resulted in reduced skin inflammation in the *Maf* KO mice compared to the WT mice (Figure 6G). Moreover, histological analysis of the dorsal skin of *Maf*-deficient mice did not show epidermal thickening pathology, a characteristic feature of psoriasis (Figures 6H and 6I). Reverse transcription polymerase chain reaction (RT-PCR) analysis of the IMQ-treated dorsal skin of *Maf* KO at day 7 revealed an absence of IL-17a mRNA compared to the treated WT (Figure 6J). Accordingly, the dorsal skin lacked ROR*γ* T^+^ *γδ* T cells in the *Maf* KO mice (Figure 6K). Taken together, *Maf*-deficient mice lack ROR*γ* T^+^ *γδ*T17 cells in the lymph nodes and epithelial tissues such as skin and intestine and are protected from IMQ-induced psoriasis-like skin inflammation.

### Sequential activation of *Sox13*, *Maf* and *Rorc* regulates the differentiation of various *γδ* T cell subsets

Next, we sought to understand the temporal dynamics and the mutual interplay of *Sox13*, *Maf* and *Rorc* in regulating *γδ*T17 differentiation. In order to investigate the causal relationship and validate the expression kinetics predicted by the pseudo-temporal ordering (i.e. *Sox13*-*Maf*-*Rorc*), we further looked into *Sox13* and *Rorc* KO mice. Since our scRNA-seq experiments identified substantial transcriptional heterogeneity and various subtypes in *γδ* T cells from the adult mouse thymi, we questioned the role of *Sox13* being essential only for the *γδ*T17 differentiation. Therefore, we investigated the role of *Sox13* in *γδ* T cell differentiation in the context of our newly identified subsets using scRNA-seq. To this end, we sequenced immature and mature *γδ* T cell subsets from *Sox13* KO mice (Figures 7A and 7B). Indeed, the immature *γδ* T cell compartment from *Sox13* KO mice lacked *Maf*^+^ *Rorc*^+^ *γδ* T cells indicating that *Sox13* acts upstream of *Maf* and *Rorc* to activate the *γδ*T17-differentiaton program (Figures 7C and 7D). Importantly, the newly identified *Gzma*^hi^ *γδ* T cell subset was also absent in these KO mice (Figures 7C and 7D). In the mature *γδ* T cell compartment, there was complete absence of *Rorc*^+^ (*Zbtb16*^+^ and *Zbtb16*^-^) *γδ* T cell subsets (Figures 7C and 7D). Interestingly, we found a large number of *Zbtb16*^+^ *γδ* T cells, which expressed neither *Rorc* nor *Tbx21* (not shown) in the WT littermates (Figure 7D). Some of these cells also expressed *Il4*, closely resembling NKT-like V*γ*1^+^ Vγ6.3^+^ *γδ* T cells (Kreslavsky et al., 2009) (Figure S4D). Of note, *Sox13* KO and WT littermates are males and were bred in a different animal facility. The number of these cells was significantly lower in the female WT data from our animal facility (Figure 2). Vice versa, the number of mature *Rorc*^+^ (*Zbtb16*^+^ and *Zbtb16*^-^) cells was much higher in our WT data (Figure 2). Nevertheless, we also observed the absence of *Zbtb16*^+^ NKT-like *γδ* T cells in the *Sox13* KO mice (Figures 7C and 7D). Differential gene expression analysis between the immature *γδ* T cells from KO and WT mice revealed downregulation of *Gzma*, *Rorc*, *Il17re*, *Blk* and *Maf* in the immature KO cells compared to the WT (Figure 7E). Interestingly, *Sox13*-deficient immature *γδ* T cells expressed higher levels of *Cd28* and *Btla*, a receptor for B7 homolog, B7x (Watanabe et al., 2003). In the mature compartment, *Rorc*, *Maf*, *Blk*, *Icos*, *Il4* and *Zbtb16* were downregulated in the KO compared to the WT (Figure 7F). In conclusion, we show that SOX13 acts upstream of c-MAF, ROR*γ* T and PLZF and regulates the differentiation of *γδ*T17 as well as NKT-like IL-4^+^ *γδ* T cells. Furthermore, it is also essential for *Gzma*^hi^ *γδ* T cell differentiation.

**Figure 7.**
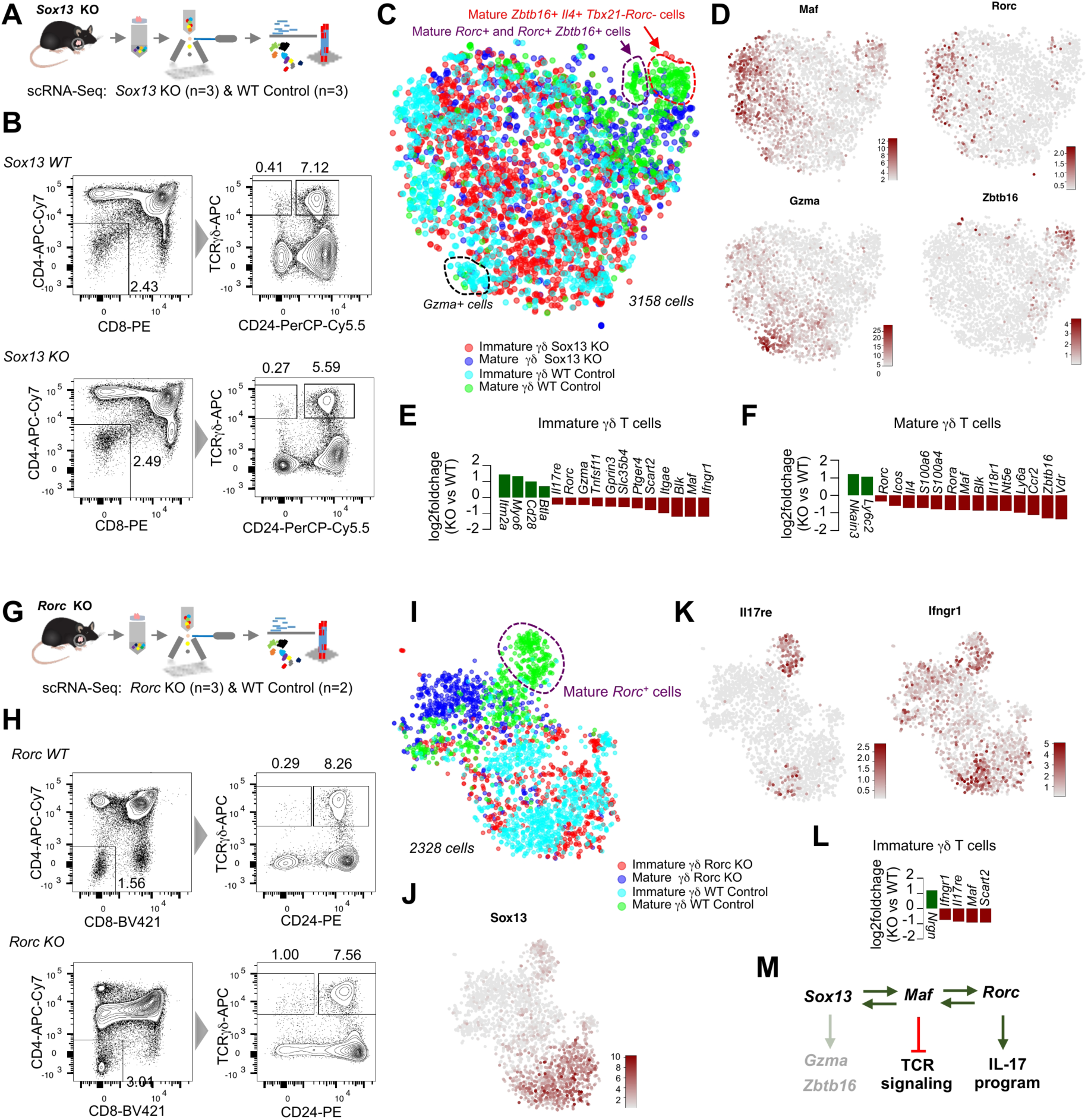
Sequential activation of *Sox13*, *Maf* and *Rorc* regulates the differentiation of various *γδ* T cell subsets. **(A)**Scheme showing the experimental design and scRNA-seq pipeline for the analysis of *Sox13* KO animals. **(B)**FACS plots depicting the sorting strategy for scRNA-seq experiments. Immature and mature *γδ* T cells were isolated from 6-week old *Sox13* KO and WT male mice (n=3, each genotype). **(C)**t-SNE representation of cell types sorted as shown in (B). Mature *Rorc*^+^ (purple) as well as *Zbtb16*^+^ (red) *γδ* T cells were absent in the *Sox13* KO mice. **(D)**t-SNE representation of the expression of *Maf*, *Rorc*, *Gzma* and *Zbtb16* for cells sorted as shown in (B). Note the absence of *Gzma*^hi^ population in the *Sox13* KO mice (outlined in blue). **(E)**Bar plot depicting the differentially expressed genes in immature *γδ* T cells between the *Sox13* KO and WT. (green: upregulated genes, red: downregulated genes, adjusted *P* < 0.05). **(F)**Bar plot depicting the differentially expressed genes in mature *γδ* T cells between the *Sox13* KO and WT. (green: upregulated genes, red: downregulated genes, adjusted *P* < 0.05). **(G)**Scheme showing the experimental design and scRNA-seq pipeline for the analysis of *Rorc* KO animals. **(H)**FACS plots depicting the sorting strategy for scRNA-seq experiments. Immature and mature *γδ* T cells were isolated from 6-week old female *Rorc* KO and WT (n=2, WT and n=3, KO). **(I)**t-SNE representation of cell types sorted as shown in (H). Note that the mature *Rorc*^+^ *γδ* T cell compartment lacks cells from the *Rorc* KO mice (outlined in purple). **(J)**t-SNE representation of the expression of *Sox13* for cell sorted as shown in (H). Note that *Sox13* expression remains unaffected in *Rorc* KO mice. **(K)**t-SNE representation of the expression of *Il17re* and *Ifngr1* for cell sorted as shown in (H). **(L)**Bar plot depicting the differentially expressed genes between immature *γδ* T cells between the *Rorc* KO and WT. (green: upregulated genes, red: downregulated genes, adjusted *P* < 0.05). **(M)**Schematic showing the regulatory hierarchy of *Sox13*, *Maf* and *Rorc* during *γδ*T17 differentiation derived from scRNA-seq analysis of KO mice for these three factors. Note that *Sox13* is also required for the development of *Gzma*^hi^ and *Zbtb16*^+^ *γδ* T cells.

Finally, we analyzed the effect of *Rorc* deficiency on *γδ* T cell differentiation. Although, like SOX13, ROR*γ* T has been shown to be essential for *γδ*T17 differentiation, its role in *γδ* T cell development in the thymus has not yet been investigated in detail. Our computational analysis revealed the activation of *Rorc* as the endpoint of the *γδ*T17 effector program, prompting us to look further into *Rorc* KO thymi. Therefore, we profiled immature and mature *γδ* T cells from *Rorc* KO thymi (Figures 7G and 7H). scRNA-seq results identified that immature *γδ* T cells from *Rorc* KO mice did not activate the *γδ*T17-differentiation program and lacked *Il17re* expression although the expression of *Sox13* remain unaffected in these cells (Figures 7I-7K). Furthermore, these cells exhibited lower expression of genes associated with *γδ*T17 differentiation such as *Maf* and *Scart2* (Figure 7L). The mature *γδ* T cell compartment completely lacked *γδ*T17 cells in the *Rorc* KO mice (Figure 7I). Taken together, we revealed that SOX13 acts as an upstream regulator of *γδ*T17 lineage specification and is essential for c-MAF-driven activation of ROR*γ* T, a transcription factor essential for *γδ*T17 differentiation (Figure 7M).

## DISCUSSION

The technical advancement in characterizing the transcriptome of single cells is enabling us to answer many fundamental questions in biology. The *γδ* T cell lineage has sparked a significant interest among immunologists because of its unique developmental paradigm and functional properties, which can be exploited for immunotherapy. scRNA-seq provided us a much needed tool to investigate their differentiation in the thymus, since progress in elucidating the factors controlling differentiation of rare *γδ* T cells in the adult thymus has been hampered by the lack of high-resolution experimental techniques (Figure 8A).

**Figure 8.**
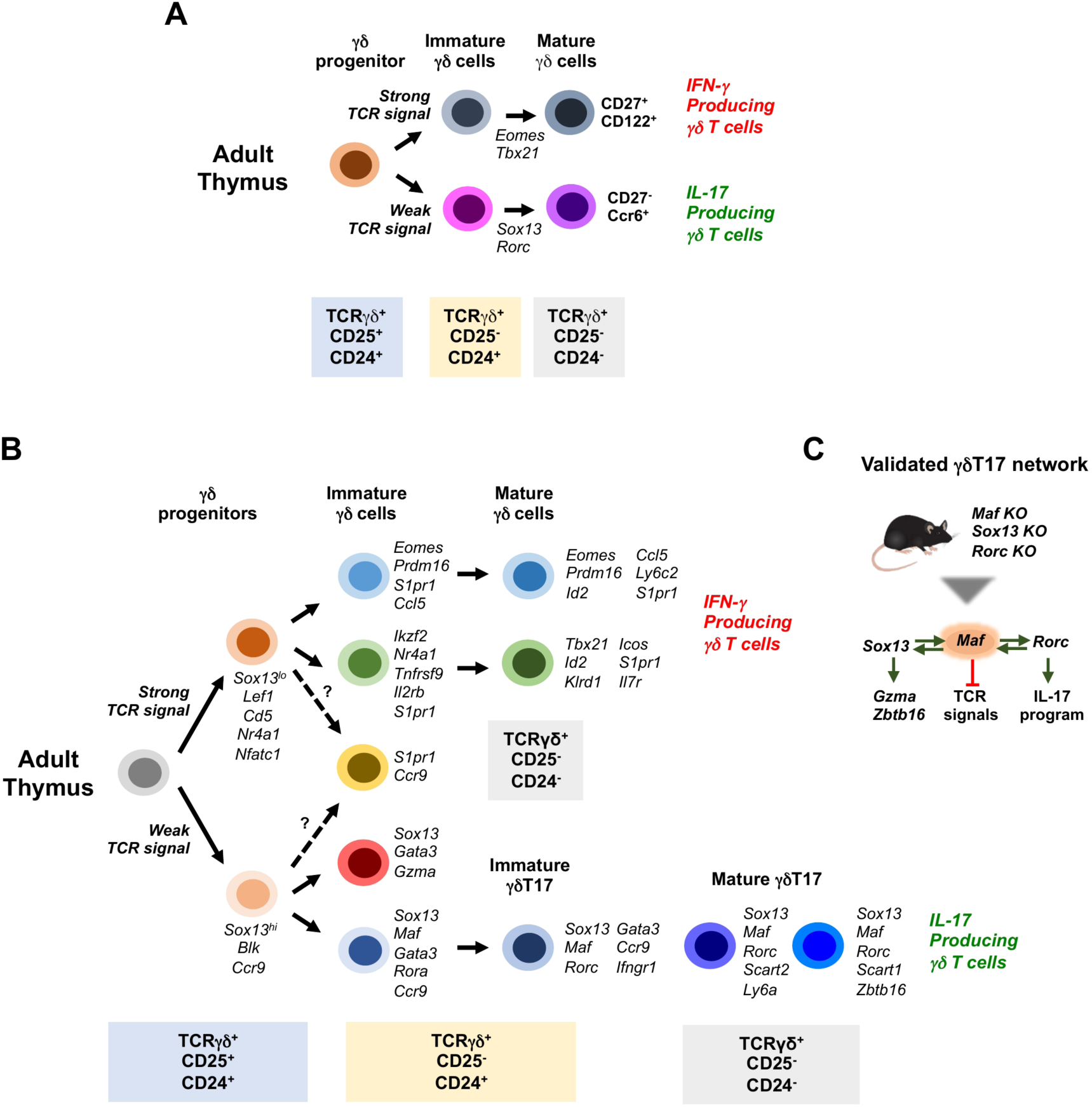
scRNA-seq of *γδ* T cell differentiation significantly enhances the understanding of their development. **(A)**Schematic showing the previously known cell types, surface markers and transcription factors characterizing *γδ* T cell differentiation. **(B)**Schematic showing the summary of *γδ* T cell differentiation revealed by scRNA-seq. We found TCR signaling-related early priming among CD25^+^ *γδ*progenitors towards *γδ*T17 and IFN-*γ* producing *γδ* T cell differentiation. We identified various novel subtypes as well as genes regulating their development. **(C)**Single-cell trajectory analysis identified *Maf* (circled in orange) as a key regulator of *γδ*T17 differentiation. We validated the role a core regulatory network comprising *Sox13*, *Maf* and *Rorc* during this differentiation process using scRNA-seq of KO mice for these three transcription factors.

The strength of TCR signals is a known determinant of *αβ*versus *γδ*lineage decision as well as *γδ* T cell effector differentiation (Haks et al., 2005; Hayes et al., 2005; Zarin et al., 2014). Therefore, it is likely that the earliest CD25^+^ *γδ*-committed progenitors at the DN3 stage experience varying levels of TCR signals that prime them towards different effector cell fates. However, due to the rarity of these cells, such transcriptional studies were never feasible. Indeed, our fate bias quantification reveals TCR signal strength-based effector lineage priming in these earliest *γδ*-committed progenitors.

A previous study has suggested SOX13 as a commitment factor influencing the *αβ*versus *γδ*lineage decision at the DN2 stage prior to TCR rearrangements (Melichar et al., 2007). These data suggested that a DN2 subset biased toward the *γδ*lineage expressed higher levels of *Sox13* than the *αβ*lineage–biased DN2 subset. Consistent with this study, we also detected low expression of *Sox13* in few cells at the DN2 stage. A differential gene expression analysis of *Sox13*^+^ versus *Sox13*^-^ cells at the DN2 stage revealed an upregulation of *Tcrg-C4*, *Gzma* and *Scart2* transcripts (data not shown), suggesting an early commitment at the DN2 stage towards the most naïve cytotoxic-like *Gzma*^hi^ population. Subsequently, TCR signaling strength could determine the fate decision towards *γδ*T17, IFN-*γ*and NKT-like lineages, respectively (Zarin et al., 2015).

Leveraging the power of scRNA-seq for rare cell type identification we discovered two previously unknown sub-types in the immature *γδ* T cell compartment of adult thymi – a *Gzma^hi^* and an *S1pr1*^+^ *Ccr9*^+^ double positive population (Figure 8B). The mature cells were more heterogeneous and revealed complex transcriptional heterogeneity (Figure 8B). Broadly, we found two subtypes in both *γδ*T17 (*Rorc*^+^ and *Rorc*^+^ *Zbtb16*^+^) and IFN-*γ* producing (*Tbx21*^+^ and *Eomes*^+^) compartments, and detected few cells co-expressing *Tbx21* and *Eomes*. Sequencing larger number of mature *γδ* T cells will help to better resolve the heterogeneity of this compartment in the future. The *γδ* T cells from the embryonic thymi were more homogeneous. Most strikingly, unlike in the adult, *Sox13*^+^ cells expressed higher levels of *Nr4a1* suggesting different mechanisms governing prenatal and postnatal *γδ* T cell differentiation.

In this study, we focused our attention to understand the regulatory mechanisms governing *γδ*T17 differentiation. Lineage tree inference and GRN analysis predicted the role of three transcription factors – *Sox13*, *Maf* and *Rorc* – in sequentially driving the differentiation of *γδ* T cell progenitors towards this lineage. Indeed, we found that *Sox13* is the earliest lineage-determining factor required for the differentiation of *Rorc*^+^, *Gzma^hi^* and *Zbtb16*^+^ *γδ* T cell subsets. In this study, we provide a detailed characterization of c-MAF, a previously unknown critical regulator of this sublineage activated downstream of SOX13. c-MAF is required for the emergence of all ROR*γ* T^+^ lineages during *γδ* T cell differentiation in the embryonic as well as the adult thymi. The sequential activation of these factors was inferred by a scRNA-seq analysis of *γδ* T cells isolated from adult thymi of knockout mice for these three factors, permitting the inference of causality. Importantly, we noticed that *Maf*-deleted cells upregulated signature genes associated with high TCR signal strength pointing to the fact that c-MAF plays an active role in modulating TCR signaling during *γδ*T17 differentiation. Interestingly, we did not see an enrichment of AKT, MAP Kinase, TLR and RTK signaling pathway genes in the *Sox13*-deficient *γδ* T cells, suggesting non-redundant roles of SOX13 and c-MAF during *γδ*T17 differentiation.

Although our study establishes a sequential and essential role of SOX13 and c-MAF in driving the differentiation of ROR*γ* T^+^ *γδ*T17 cells (Figure 8C), the precise mechanistic details of the regulatory interactions among these three transcription factors remain to be investigated in future studies. Previously, it has been shown that SOX5 interacts with the DNA-binding domain of c-MAF via its HMG domain and cooperatively activates the promoter of ROR*γ* T in CD4^+^ T cells during T helper 17 differentiation (Tanaka et al., 2014). We therefore suspect a similar regulatory landscape where SOX13/SOX4 physically interact with c-MAF to activate the ROR*γ* T promoter, leading to the activation of *γδ*T17 specific differentiation program during *γδ* T cell differentiation. Furthermore, since ROR*γ* T is also required for *αβ* T cell development in the thymus, we anticipate differential chromatin regulatory mechanisms driving its expression during the differentiation of *αβ*and *γδ* T cell lineages. Chromatin accessibility and immunoprecipitation assays on the purified *γδ* T cell subsets will be essential to dissect the intricacies of regulatory cascades driving the *γδ* T cell differentiation program.

In summary, our work reveals continuous high-resolution intrathymic differentiation trajectories of the *γδ* T cell lineages. Our results substantially increase the knowledge about the development of this understudied rare branch of T cells at the interface of the adaptive and the innate immune system, which has lately received increasing attention as crucial player in autoimmune diseases and cancer. Here, we could comprehensively delineate progenitor stages of the *γδ* T cell differentiation trajectories, identify and validate the role of a novel critical transcription factor as well as infer the order of regulatory events leading to the emergence *γδ*T17 cells in the adult thymus (Figures 8B and 8C).

More broadly, our study serves as a proof-of-principle that a powerful combination of flow cytometry and scRNA-seq in wild-type and knockout animals of candidate regulators combined with state-of-the-art computational analysis can provide the tools to decipher differentiation trajectories even of rare cell types.

## DATA AVAILABILITY

The primary read files as well as expression count files for the single-cell RNA-sequencing datasets reported in this paper are available to download from GEO (accession number: GSE115765).

## AUTHOR CONTRIBUTIONS

D.G. conceived the study, Sagar and D.G. designed the experiments, Sagar performed all FACS and scRNA-seq experiments as well as data analysis under the supervision of D.G., M.P. performed thymocyte isolation from *Maf* KO mice and characterized the phenotype of *Maf* KO mice, J.S.H. contributed to FACS and scRNA-seq experiments, M.P. and S.N. performed the psoriasis experiments, E.S. and U.L. contributed to *Sox13* and *Rorc* KO mice work respectively, M.W., Y.T. and D.R.L. supervised the *Sox13*, *Rorc* and *Maf* KO studies respectively. Sagar wrote the manuscript with guidance by D.G. and input from M.P. and D.R.L. All authors edited the manuscript.

## ACKNOWLEDGMENTS

The authors would like to thank Thomas Boehm for his valuable support during the course of study and reading the manuscript; Christiane Happe and Ingrid Falk for their assistance with the mouse work and flow cytometry; Patrice Zeis and Anja Nusser for critical reading of the manuscript; Sebastian Hobitz, Konrad Schuldes, Maike Knoblauch and Andreas Würch from the flow cytometry facility as well as Ulrike Bönisch and Laura Arrigoni from the deep sequencing facility. This work was supported by DFG grant GR4980/3-, the Behrens-Weise-Foundation, and by the Max Planck Society.

## METHODS

### Mice

C57BL/6J mice were purchased from Charles River or obtained from in-house breeding. Mice were kept in the animal facility of the Max Planck Institute of Immunobiology and Epigenetics in specific-pathogen-free (SPF) conditions. All animal experiments were performed in accordance with the relevant guidelines and regulations, approved by the review committee of the Max Planck Institute of Immunobiology and Epigenetics and the Regierungspräsidium Freiburg, Germany. Generation and genotyping of *Sox13*-deficient (*Sox13* KO) mice has been described previously (Baroti et al., 2016). *Sox13* KO mice were on a C57BL/6J background. Experiments were approved by the responsible local committees and government bodies (University, Veterinäramt Stadt Erlangen & Regierung von Unterfranken). *Maf* KO mice were bred and maintained in the animal facility of the Skirball Institute (New York University School of Medicine) in SPF conditions. C57Bl/6 mice were obtained from Jackson Laboratories or Taconic Farm. *Maf^fl/fl^* and *Il7ra^Cre^* mice were kindly provided by C. Birchmeier and H. R. Rodewald (Schlenner et al., 2010; Wende et al., 2012). *Maf* conditional KO mice were generated by crossing *Maf^fl/fl^* to *Il7ra^Cre^* animals. All animal procedures were performed in accordance with protocols approved by the Institutional Animal Care and Usage Committee of New York University School of Medicine. Generation of *Rorc* KO (*Rorc^tm2Litt^*) mice has been described previously (Eberl et al., 2004). *Rorc^tm2Litt^* mice were bred and maintained in the animal facility of Institute of Medical Microbiology and Hygiene at University Medical Center Freiburg and experiments were approved and are in accordance with the local animal care committees (Regierungspräsidium Freiburg).

### Thymocyte isolation

All animals were sacrificed using carbon dioxide and cervical dislocation. To isolate thymocytes, thymus was dissected and placed on a 40 µm cell strainer (Falcon, Corning) kept on a 50 ml tube (Falcon, Corning). Thymus was mashed on the cell strainer using the back of the 1 ml syringe plunger. 10 ml Phosphate-buffered saline (PBS) was continuously added while mashing the thymus to collect the single cell suspension in the 50 ml tube. Collected thymocytes were centrifuged at 400*g* for 5 min at 4ºC. The pellet was dissolved in 10 ml PBS and passed through the 30 µm nylon filter (CellTrics, Sysmex) kept on a 15 ml tube (Falcon, Corning). Cells were again centrifuged at 400*g* for 5 min at 4ºC. Afterwards the pellet was dissolved in 200 µl of PBS.

### Magnetic enrichment of double negative thymocytes

After thymocyte isolation as describe above, the pellet was dissolved in 3 ml of PBS. 3 ml of the following biotin-labeled antibody cocktail solution (1:50 dilution each, clones are mentioned in the barckets) was prepared: CD8a (53-6.7), CD4 (GK1.5), NK-1.1 (PK136), TER-119 (TER-119), Ly-6G/Ly-6C

(RB6-8C5) and CD11c (N418). All antibodies were purchased from BioLegend. The antibody solution was incubated with the thymocytes for 15 min on ice. The cells were centrifuged at 400*g* for 5 min at 4ºC and washed with 5 ml of PBS. The resulting pellet was re-suspended in 600 µl of PBS and 100 µl streptavidin-conjugated beads (MojoSort Streptavidin Nanobeads, Biolegend) were added to the solution and incubated for 15 min on ice. Afterwards, the tube was placed on the magnet for 5 min to let the beads firmly attached to the wall of the tube in contact with the magnet. The remaining liquid containing the cells without the beads was transferred into a 1.5 ml tube (Eppendorf) and centrifuged at 400*g* for 5 min at 4ºC. The pellet was dissolved in 50 µl of PBS.

### Cryopreservation and thawing of thymocytes

Thymocytes from *Maf* KO and *Sox13* KO mice as well as the corresponding littermate controls were cryopreserved prior to shipment on dry ice to the Max Planck Institute of Immunobiology and Epigenetics, Freiburg. Thymocytes were isolated as described before and resuspended in 4 ml of freezing medium containing 90 % fetal bovine serum (FBS) and 10 % Dimethyl sulfoxide (DMSO) and kept at - 80ºC overnight before shipment. After receiving the frozen thymocytes, they were kept at - 80ºC for few days. Prior to antibody staining, frozen cells (in 90% FBS and 10% DMSO) were de-frozen in the water bath at 37°C until almost thawed. The cells were transferred to a 15 ml tube (Falcon, Corning) and 14 ml of pre-warmed RPMI medium 1640 (Gibco) was added drop-wise while gently swirling and inverting the tube. The cells were then centrifuged at 400*g* for 5 min at 4°C. The supernatant was discarded and cells were washed again with 10 ml RPMI medium 1640. After centrifugation, the cell pellet was re-suspended in 200 μl of RPMI medium 1640.

### Antibody staining, flow cytometry and single cell sorting

200 µl of antibody staining solution was prepared in PBS (for freshly isolated thymocytes) or RPMI medium 1640 (for frozen lymphocytes) and added to the 200 μl of re-suspended thymocyte solution. For enriched double negative thymocytes, 50 µl of antibody solution was added to the pellet, which was dissolved in 50 µl of PBS. Afterwards, thymocytes in the antibody solution were incubated for 20 min on ice. Cells were then washed twice with 1 ml of PBS or RPMI medium 1640 and re-suspended in 3 ml after the last wash. Just prior to single cell sorting using a flow cytometer, 5 μl of 20 μg/ml 4′,6-diamidino-2-phenylindole (DAPI, Sigma) solution was added to the tube to stain for dead cells. Flow cytometry data was analyzed using the FlowJo program. The following antibodies were used (clones and used dilutions are mentioned in the brackets): CD117-BV510 (ACK2, 1:500), CD44-PerCP/Cy5.5 (IM7, 1:500), CD25-BV421 (PC61, 1:500), CD122-PE (TM-β1, 1:500), CD8a-BV421 (53-6.7, 1:1000), CD8a-FITC (53-6.7, 1:500) CD4-PE (RM4-5, 1:1000), CD4-APC/Cy7 (RM4-5, 1:500), CD4-APC (RM4-5, 1:500), TCR*γδ*-APC (GL3, 1:500), CD24-PE (M1/69, 1:1000), CD24-PerCP/Cy5.5 (M1/69,1:500), V*γ*1.1 (2.11, 1:500), V*γ*2-FITC (UC3-10A6, 1:500), V*γ*3-PE (536, 1:500) and V*γ*6.3/2-PE (8F4H7B7, 1:500). All the antibodies were purchased from BioLegend except CD8a-FITC, CD44-PerCP/Cy5.5 and V*γ*6.3/2-PE (BD Pharmingen). Single cells were sorted in 384-well plates (Bio-Rad Laboratories) containing lysis buffer and mineral oil (see the next section) using BD FACSAria FUSION. The sorter was run on single cell sort mode. Using pulse geometry gates (FSC-W x FSC-H and SSC-W x SSC-H), doublets were excluded. After the completion of sorting, the plates were centrifuged for 10 min at 2200*g* at 4°C, snap-frozen in liquid nitrogen and stored at -80°C until processed.

### Single-cell RNA amplification and library preparation

Single-cell RNA sequencing was performed using the mCEL-Seq2 protocol, an automated and miniaturized version of CEL-Seq2 on a mosquito nanoliter-scale liquid-handling robot (TTP LabTech) (Hashimshony et al., 2016; Herman et al., 2018). Twelve libraries with 96 cells each were sequenced per lane on Illumina HiSeq 2500 or 3000 sequencing system (pair-end multiplexing run) at a depth of ~130,000-200,000 reads per cell.

### Quantification of transcript abundance

Paired end reads were aligned to the transcriptome using bwa (version 0.6.2-r126) with default parameters (Li and Durbin, 2010). The transcriptome contained all gene models based on the mouse ENCODE VM9 release downloaded from the UCSC genome browser comprising 57,207 isoforms, with 57,114 isoforms mapping to fully annotated chromosomes (1 to 19, X, Y, M). All isoforms of the same gene were merged to a single gene locus. Furthermore, gene loci overlapping by >75% were merged to larger gene groups. This procedure resulted in 34,111 gene groups. The right mate of each read pair was mapped to the ensemble of all gene loci and to the set of 92 ERCC spike-ins in sense direction (Baker et al., 2005). Reads mapping to multiple loci were discarded. The left read contains the barcode information: the first six bases corresponded to the unique molecular identifier (UMI) followed by six bases representing the cell specific barcode. The remainder of the left read contains a polyT stretch. For each cell barcode, the number of UMIs per transcript was counted and aggregated across all transcripts derived from the same gene locus. Based on binomial statistics, the number of observed UMIs was converted into transcript counts (Grun et al., 2014).

### Clustering and visualization

Clustering analysis and visualization were performed using the RaceID3 algorithm (Herman et al., 2018). The number of quantified genes ranged from 25,718 to 26,970. Cells with a total number of transcripts <2,500 were discarded and count data of the remaining cells were normalized by downscaling. Cells expressing >2% of *Kcnq1ot1*, a potential marker for low-quality cells (Grun et al., 2016), were not considered for analysis. Additionally, transcript correlating to *Kcnq1ot1* with a Pearson’s correlation coefficient >0.65 were removed. The following parameters were used for RaceID3 analysis: mintotal=2500, minexpr=5, outminc=5, FSelect=TRUE, probthr=10^−4^. CGenes was initialized with the following set of genes to remove cell-cycle-associated and batch-associated variability: *Pcna*, *Mki67*, *Mir703*, *Gm44044*, *Gm22757*, *Gm4775*, *Gm17541*, *Gm8225*, *Gm8730*, *Ptma*, *Actb*, *Hsp90aa1*, *Hsp90ab1* and *Ppia*. Reclassification of cells based on random forests was performed after outlier identification.

### Differential gene expression analysis

Differential gene expression analysis was performed using the diffexpnb function of RaceID3 algorithm. Differentially expressed genes between two subgroups of cells were identified similar to a previously published method (Anders and Huber, 2010). First, negative binomial distributions reflecting the gene expression variability within each subgroup were inferred based on the background model for the expected transcript count variability computed by RaceID3. Using these distributions, a p value for the observed difference in transcript counts between the two subgroups was calculated and multiple testing corrected by the Benjamini-Hochberg method.

### FateID analysis

In order to investigate the transcriptional priming in early CD25^+^ *γδ* T cell progenitors, FateID (Herman et al., 2018) was run on clusters 6, 11, 13, 17 and 18 as well as TCR*γδ*^+^ cells from clusters 12 and 14 with minnr=5 and minnrh=10. Clusters 11 (IL-17-primed) and 18 (IFN-*γ*-primed) were used as target clusters. Classical multidimensional scaling was used for dimensional reduction and visualization of the results. Differential gene expression analysis was performed between cells biased towards one of the lineages with fate bias probability >0.5 using diffexpnb.

### Lineage inference and pseudo-temporal ordering

To derive the differentiation trajectories of *γδ* T cell differentiation, the StemID2 algorithm was used (Grun et al., 2016; Herman et al., 2018). StemID2 was run with the following parameters: cthr=15, pdishuf=1000, pthr=0.01, pethr=0.05, scthr = 0.6. Based on the *γδ*T17 differentiation trajectory predicted by StemID2, cells comprised of clusters 3, 2, 9, 4, 15, 16, 14, 6, 17, 12, 18, 19 and 21 were used to compute the self-organizing maps of pseudo-temporal expression profiles of the *γδ*T17 lineage. Similarly, clusters 3, 2, 9, 4, 15, 16, 14, 6, 17, 13, 11 and 8 were used to compute the self-organizing maps of pseudo-temporal expression profiles of the IFN-*γ* producing lineage.

### Gene regulatory network inference

Gene regulatory network inference was performed using the random forests-based ensemble method of GENIE3 algorithm (Huynh-Thu et al., 2010). Only genes expressed at >5 transcripts in at least one of the cells were included in the analysis. R/C implementation of GENIE3 was used for network inference. The top 1500 regulatory link with highest weights were shortlisted for visualization of the network using Cytoscape with default settings (Shannon et al., 2003).

### Gene set enrichment analysis

Gene set enrichment analysis was performed using gsePathway function of ReactomePA, an R/Bioconductor package (Yu and He, 2016). The fold change for each gene was calculated between *Maf* KO and WT immature *γδ* T cells using the diffexpnb function of RaceID3 and was given as an argument to gsePathway function to calculate enriched gene sets in KO cells using the following parameters: nPerm=1000, minGSSize=120, pvalueCutoff=0.05, pAdjustMethod="BH", organism = "mouse".

### Skin inflammation models

The dorsal skin of 8-week-old mice in the telogen (resting) phase of the hair cycle were shaved with clippers and then subjected to topical application or treatment of the skin as below. IMQ: mice were treated with either ~1 mg cm^−2^ skin of 5% IMQ cream (Perrigo) or control Vanicream (Pharmaceutical Specialties Inc.) for 6 consecutive days as previously described (van der Fits et al., 2009).

### Cell isolation, tissue processing and flow cytometry

Keratinocyte isolation was adapted from a previously described protocol (Nowak and Fuchs, 2009). In brief, dorsal skin was shaved and digested using either 0.25% trypsin/EDTA (Gibco) or collagenase (Sigma) to obtain a single-cell suspension. Immune cells from 1 cm^2^ pieces of skin were isolated after digestion with liberase (Roche) based on an adapted protocol (Keyes et al., 2016). Female mice were used for sorting experiments at all timepoints and conditions to obtain maximal cell numbers. Single-cell suspensions were stained with antibodies at predetermined concentrations in a 100 μl staining buffer (PBS containing 5% FBS and 1% HEPES) per 10^6^ cells. Stained cells were re-suspended in DAPI in FACS buffer (Sigma) before analysis. Data were acquired on LSRII Analyzers (BD Biosciences) and then analysed with FlowJo program. FACS was conducted using Aria Cell Sorters (BD Biosciences) into either staining buffer or Trizol LS (Invitrogen).

### Histology

Skin tissue was fixed in PBS containing 10% formalin, paraffin embedded, sectioned (0.8 mm) and stained with haematoxylin and eosin by Histowiz Inc. Stained slides were scanned at 40X magnification using Aperio AT2. Slides were visualized and epidermal thickness was analysed manually based on morphological features of the epidermis using the Aperio Image Scope software. Each skin section was measured at 10 different locations at least 10 mm apart and averaged to obtain presented thickness value.

### RNA purification and quantitative PCR

Individual animals were used for qPCR experiments. Total RNA was purified from either whole skin biopsies, flash frozen and then homogenized with a Bessman Tissue Pulverizer (SpectrumTM) or FACS-purified keratinocyte populations using Direct-zol RNA MiniPrep kit (Zymo Research) as per manufacturer’s instructions. Equal amounts of RNA were reverse-transcribed using the superscript VILO cDNA synthesis kit (Invitrogen). cDNAs for each sample were normalized to equal amounts using primers against *Actb*. XpressRef Universal Total RNA (Qiagen) was used as a negative control to assess FACS population purity.

### Isolation and stimulation of SILP and LILP lymphocytes

Intestinal tissues were sequentially treated with PBS containing 1 mM DTT at room temperature for 10 min, and 5 mM EDTA at 37°C for 20 min to remove epithelial cells, and then minced and dissociated in RPMI containing collagenase (1 mg/ml collagenase II; Roche), DNase I (100 µg/ml; Sigma), dispase (0.05 U/ml; Worthington) and 10% FBS with constant stirring at 37°C for 45 min (SI) or 60 min (LI). Leukocytes were collected at the interface of a 40%/80% Percoll gradient (GE Healthcare). For cytokine analysis, lamina propria mononuclear cells were incubated for 5 h in RPMI with 10% FBS, phorbol 12-myristate 13-acetate (PMA) (50 ng/ml; Sigma), ionomycin (500 ng/ml; Sigma) and GolgiStop (BD). Cells were stained for surface markers before fixation and permeabilization, and then subjected to intracellular cytokine staining according to the manufacturer’s protocol (Intracellular Fixation & Permeabilization buffer set from eBioscience).

---

**Figure S1.**
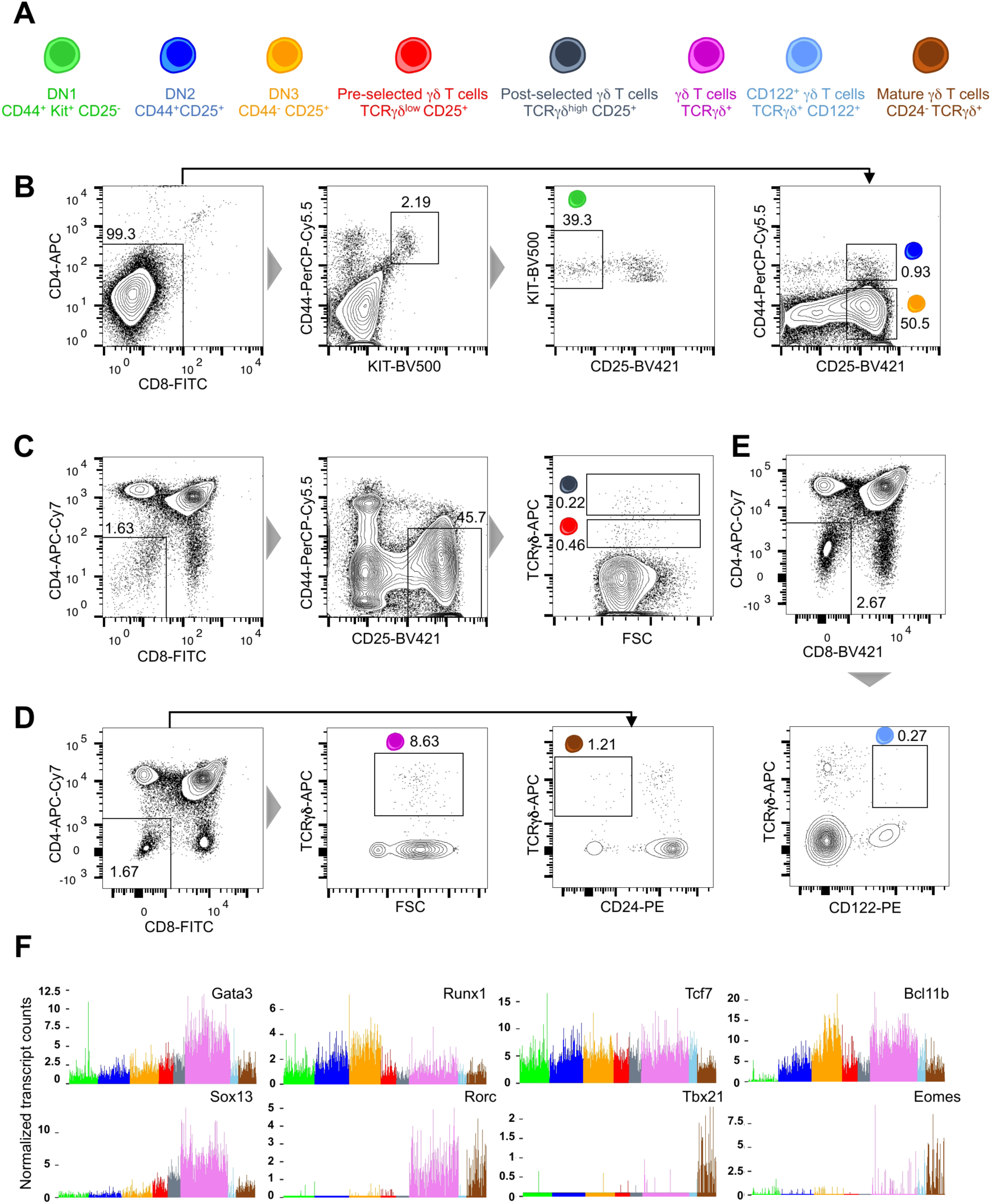
FACS strategy for scRNA-seq experiments. **(A)**Sketch showing different cell types sorted for scRNA-seq experiments and the associated cell surface markers used. **(B)**FACS plots showing the gates used for sorting DN1, DN2 and DN3 T cells. Note that before sorting DN1-DN3 populations, thymocytes were enriched for DN populations using magnetic cell enrichment. **(C)**FACS plots showing the gates used for sorting pre-selected and post-selected *γδ* T cells. **(D)**FACS plots showing the gates used for sorting pan *γδ* T cells and CD24^-^ mature *γδ* T cells. Note that >98 % of the pan *γδ* T cells are immature *γδ* T cells. **(E)**FACS plots showing the gate used for sorting CD122^+^ *γδ* T cells. **(F)**Bar plots showing the normalized transcript counts of key genes implicated in T cell commitment and *γδ* T cell differentiation. Colors of the bars represent the corresponding sorted cell types depicted in (A).

**Figure S2.**
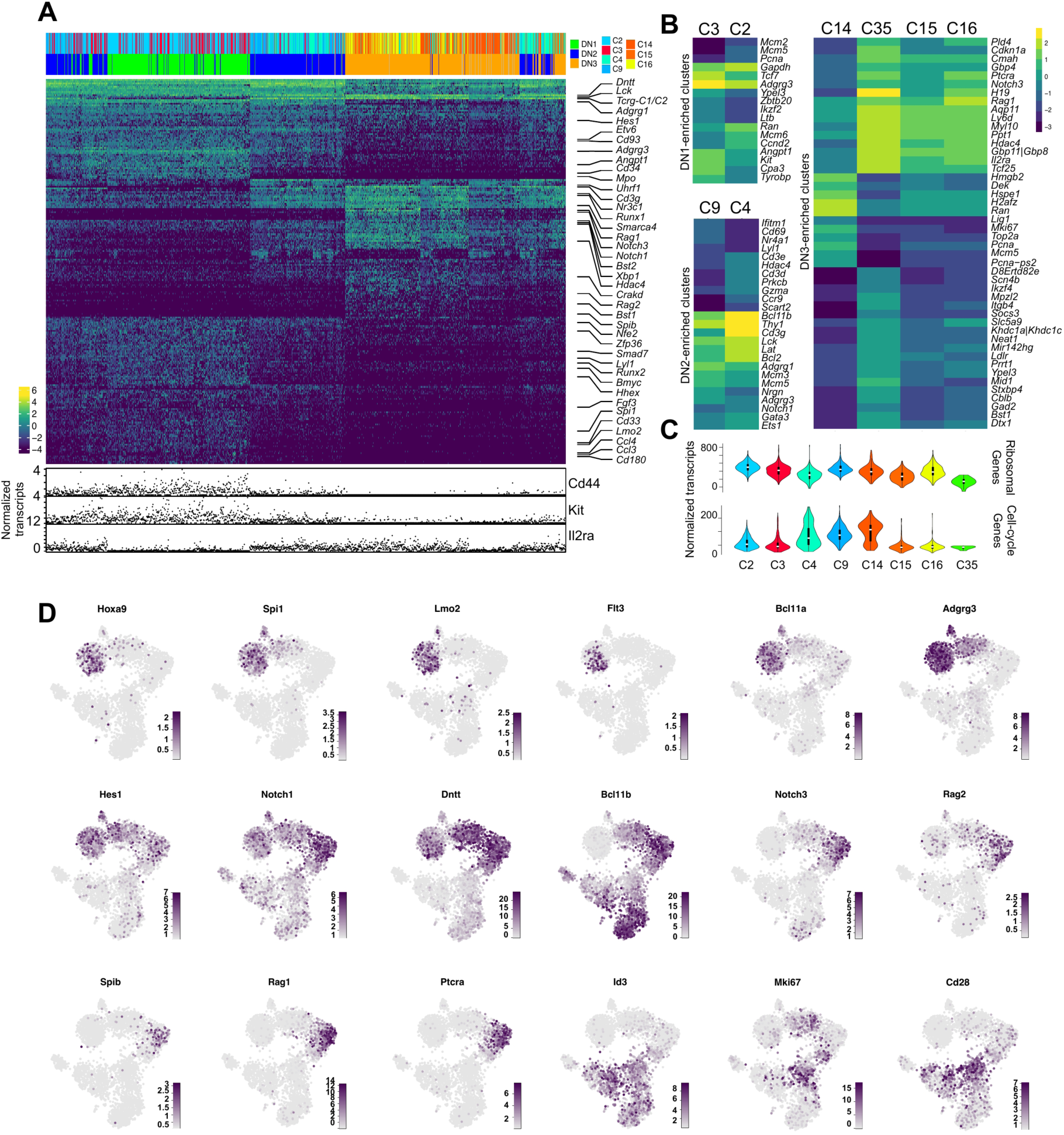
Cellular heterogeneity in the double negative T cell progenitors. **(A)**Heatmap showing the differentially expressed genes in clusters comprised of DN1-DN3 cells. Few marker genes are labeled. The normalized transcript counts for *Cd44*, *Kit* and *Il2ra*, genes distinguishing the different DN compartments, are shown below the heatmap. Only genes showing a minimum of 4-fold change in one of the pairwise cluster comparison were included (adjusted *P* < 0.05). **(B)**Heatmap showing the differentially expressed genes in DN1 (clusters 2 and 3), DN2 (clusters 9 and 4) and DN3 (clusters 15, 15, 16 and 35) clusters (adjusted *P* < 0.05). **(C)**Violin plots showing the aggregated normalized transcript counts for ribosomal and cell cycle-related genes in DN1-DN3 enriched clusters. **(D)**t-SNE representations showing the expression of key genes implicated in T cell commitment and differentiation. Scale represents normalized transcript counts.

**Figure S3.**
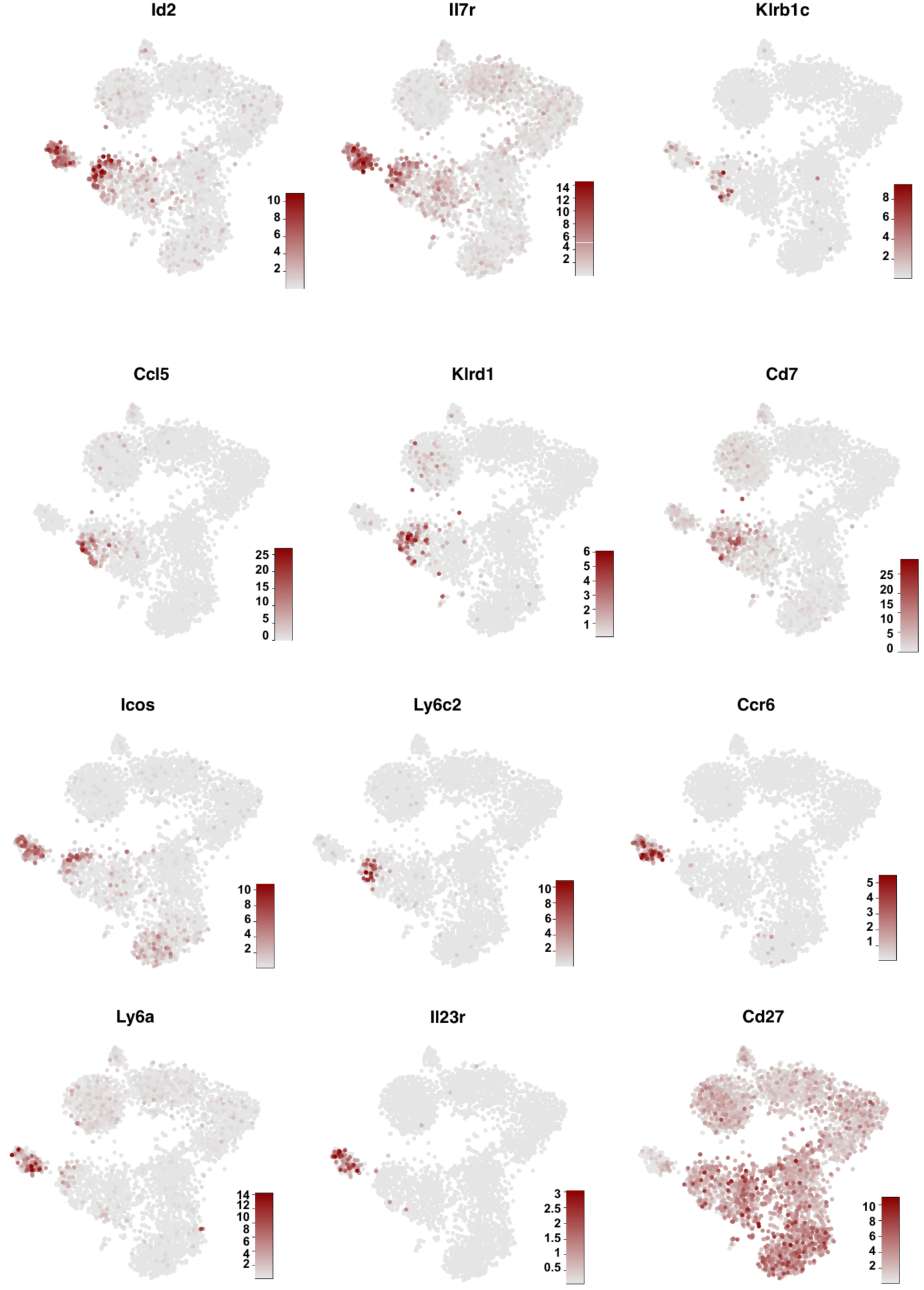
Key marker genes characterizing *γδ* T cells in the adult thymi. **(A)**t-SNE representation showing the expression of heterogeneous marker genes in immature and mature **(B)***γδ* T cell compartments in addition to those shown in Figure 2C.

**Figure S4.**
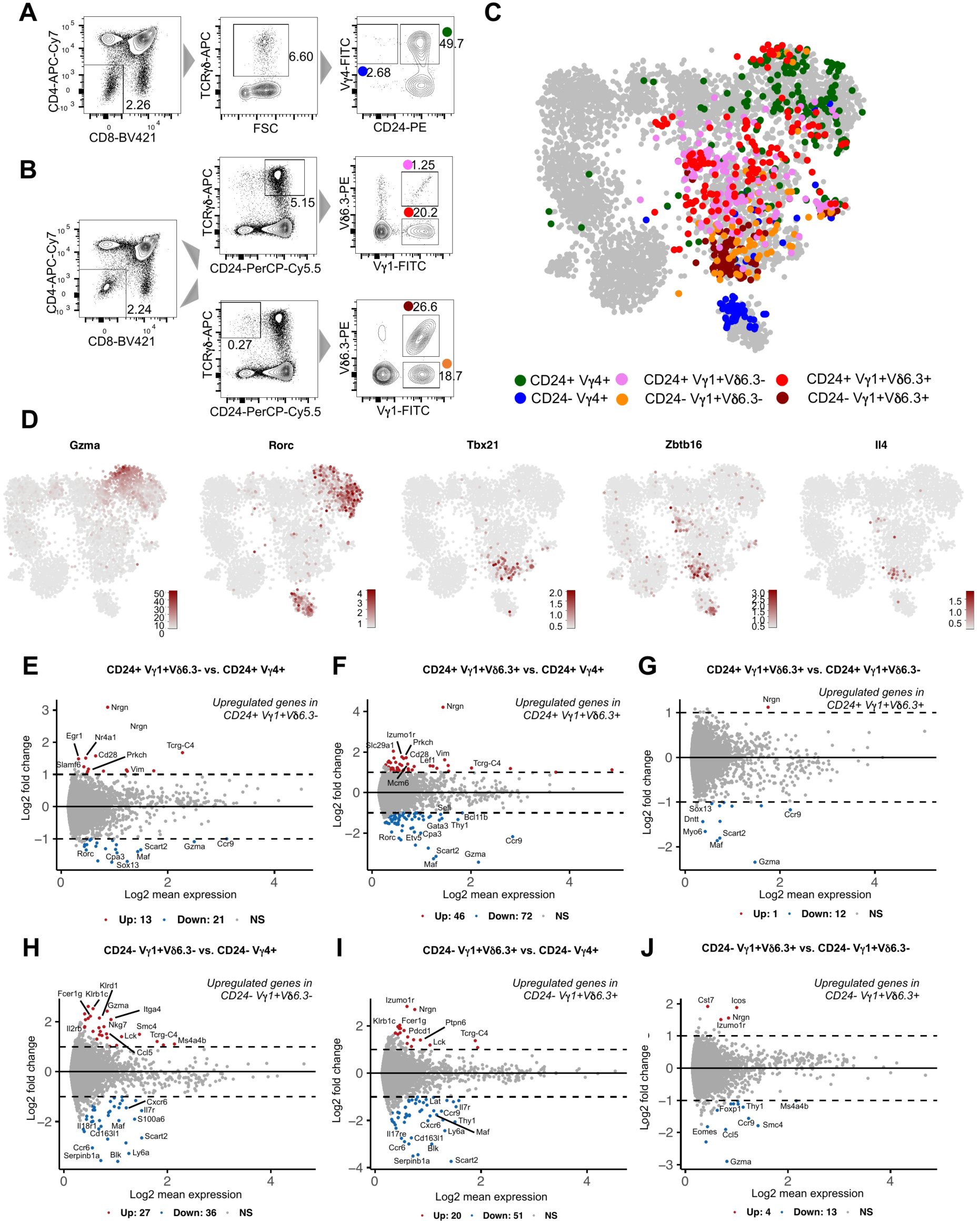
scRNA-seq of adult *γδ* T cells expressing different TCR *γ* and *δ* chains. **(A)**FACS plots showing the single cell sorting strategy for isolating immature and mature V*γ*4^+^ *γδ* T cells from 6-week old female mice (n=2). **(B)**FACS plots showing the single cell sorting strategy for isolating immature and mature V*γ*1^+^ and V*γ*1^+^V*γ*6.3^+^ *γδ* T cells (n=2, 6-week old female mice). **(C)**t-SNE representation based on transcriptome similarities showing the sorted cell types depicted in different colors except grey. t-SNE map represents all the cells including those shown in Figure 1C (shown in grey). **(D)**t-SNE representation showing the expression of *Gzma*, *Rorc*, *Tbx21*, *Zbtb16* and *Il4*. Note the expression of *Zbtb16* and *Il4* in immature and mature V*γ*1^+^V*γ*6.3^+^ *γδ* T cells. (E-J) MA plots showing the differentially expressed genes between the cells expressing different TCR*γ*and *δ* chains. Genes showing a minimum of 2-fold change (adjusted *P* < 0.05) are labeled in red (upregulated) and blue (downregulated). Names of the few key genes are highlighted.

**Figure S5.**
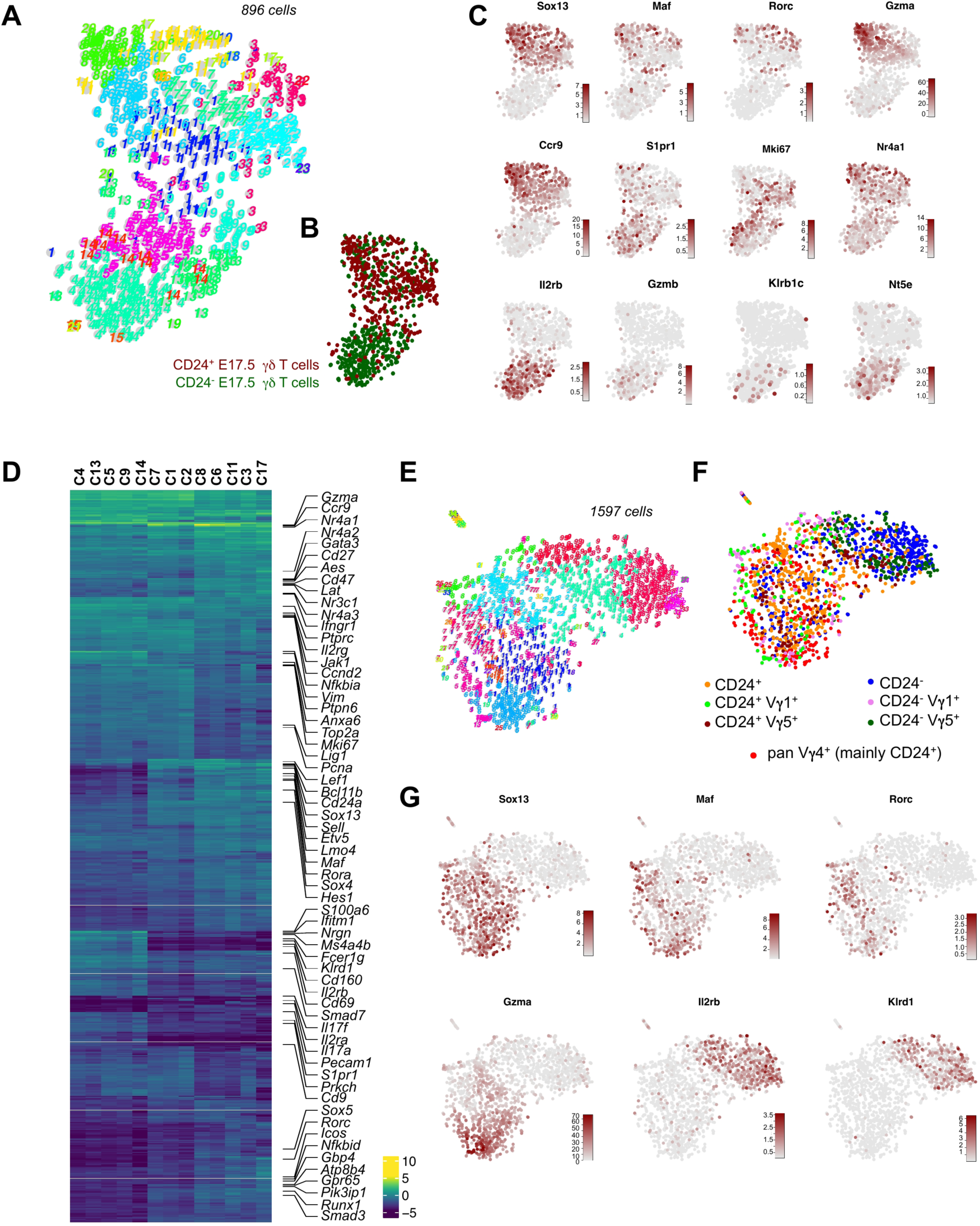
scRNA-seq analysis of *γδ* T cells from the embryonic day 17.5 thymi. **(A)**t-SNE representation of RaceID3 clusters identified in the dataset. **(B)**t-SNE maps showing the sorted cells types. Only Cd24^+^ and Cd24^-^ cells were sorted. **(C)**t-SNE representation showing the expression of key marker genes identified in the dataset. Many *γδ* T cells were proliferative as they expressed *Mki67*. **(D)**Heatmap showing the differentially expressed genes in the RaceID3 clusters. Genes showing a minimum of 2-fold change in one of the pairwise cluster comparison were included (adjusted *P* < 0.05). Few key genes are labeled. **(E)**t-SNE representation of RaceID3 clusters. Dataset comprised of *γδ*cells shown in Figure S5A as well as *γδ* T cells expressing different TCR*γ*and *σ* chains expressed on their cell surface (V*γ*1, V*γ*4 and V*γ*5). **(E)**t-SNE representation showing the different cell types sorted. **(F)**t-SNE representations showing the expression profiles of key marker genes detected in the dataset.

**Figure S6.**
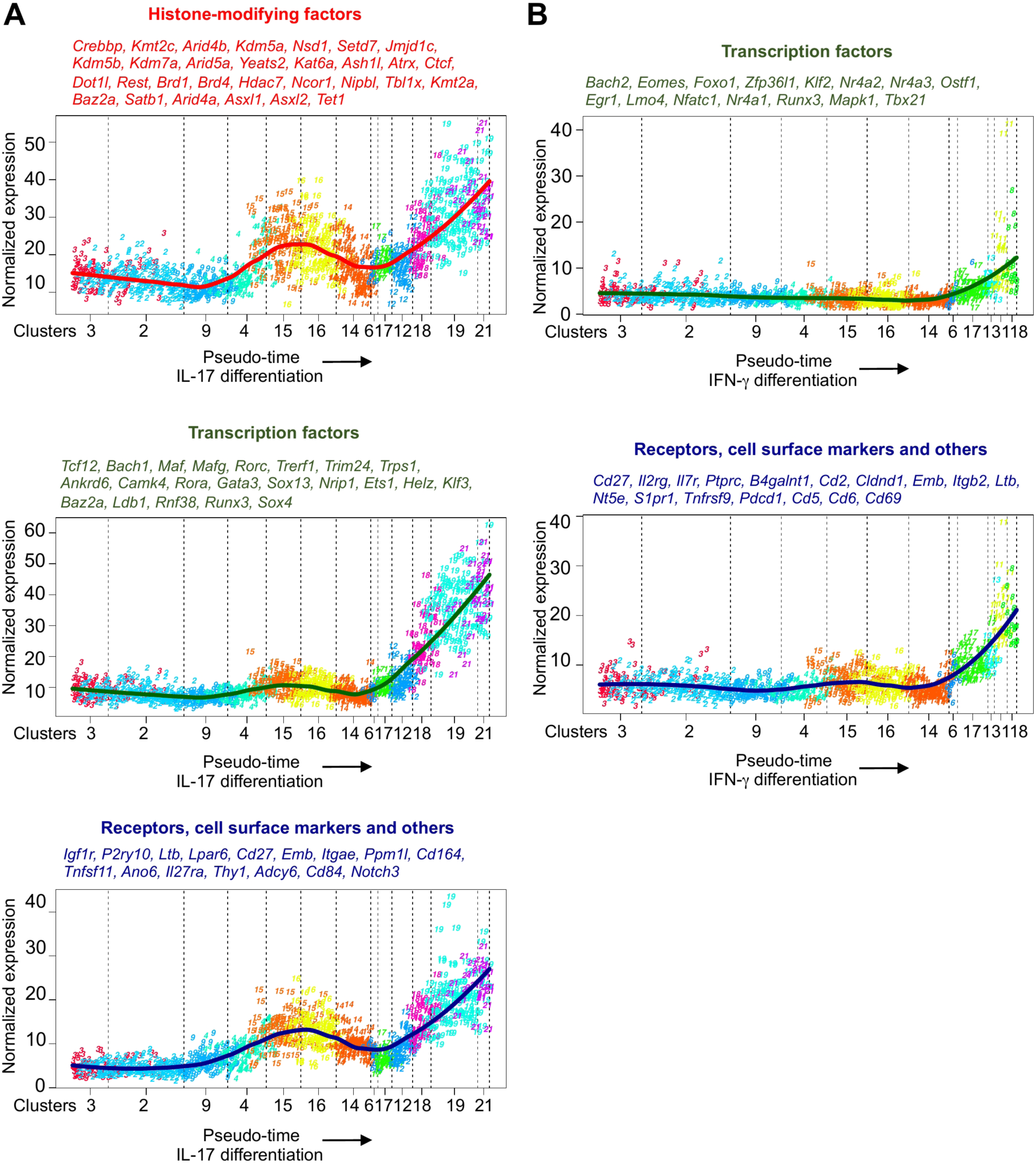
Pseudo-time analysis of *γδ*T17 and IFN-γ producing *γδ* T cell differentiation trajectories. **(A)**Pseudo-time analysis of *γδ*T17 differentiation reveals many regulators with known and unknown functions. Histone-modifying factors were specifically upregulated during *γδ*T17 differentiation. X-axis represents aggregated normalized counts of genes shown on the top of each plot. Y-axis represents the pseudo-time order shown in Figure 3B. The lines indicate the pseudo-temporal expression values derived by a local regression of expression values across the ordered cells. **(B)**Pseudo-time analysis of IFN-*γ* producing *γδ* T cell differentiation also reveals many regulators with known and unknown functions. The expression of TCR signal strength-related genes – *Nr4a1*, *Egr1*, *Cd5* and *Cd69* was upregulated during IFN-*γ* producing *γδ* T cell differentiation. X-axis represents aggregated normalized counts of genes shown on the top of each plot. Y-axis represents the pseudo-time order shown in Figure 3C. The lines indicate the pseudo-temporal expression values derived by a local regression of expression values across the ordered cells.

